# Hunger signalling in the olfactory bulb primes exploration, food-seeking and peripheral metabolism

**DOI:** 10.1101/2023.01.26.525804

**Authors:** Romana Stark, Harry Dempsey, Elizabeth Kleeman, Martina Sassi, Jeffrey Davies, Jeffrey M. Zigman, Zane B. Andrews

## Abstract

Growing evidence highlights a complex interaction between olfaction and metabolism with impaired olfactory function observed in obesity and increased olfactory sensitivity during hunger. The mechanisms linking metabolic state and olfaction remain unknown, but increased accessibility of hormones, such as ghrelin, and the diverse expression of hormone receptors such as those for ghrelin (GHSRs) in the olfactory system suggests an underappreciated neuroendocrine role. Here, we examined the hypothesis that GHSRs in the olfactory bulb (OB) link hunger with olfactory sensitivity to influence foraging behaviours and metabolism. Selective deletion of OB^GHSRs^ in adult male mice was achieved with adeno-associated viral expression of cre-recombinase in the OB of floxed-*Ghsr* mice. OB^GHSR^ deletion significantly affected olfactory discrimination and habituation to both food and pheromone odours, with greatest effect under fasted conditions. Anxiety-like and depression-like behaviour was significantly greater after OB^GHSR^ deletion using 3 independent anxiety behavioural tasks and testing for anhedonia, whereas exploratory behaviour was reduced. No effect on spatial navigation and memory was observed. Although OB^GHSR^ deletion did not affect cumulative food intake, it significantly impacted feeding behaviour as evidenced by altered bout number and duration. Moreover, food-finding after fasting or ip ghrelin was attenuated. Intriguingly, OB^GHSR^ deletion caused an increase in body weight and fat mass, spared fat utilisation on a chow diet and impaired glucose metabolism indicating metabolic dysfunction. We conclude that OB^GHSRs^ maintain olfactory sensitivity, particularly during hunger, and facilitate behavioural adaptations that optimise food-seeking in anxiogenic environments, priming metabolic pathways in preparation for food consumption.

## Introduction

The neural control of appetite and metabolism plays an essential role in the maintenance of body weight and dysfunction in these circuits may exacerbate metabolic disease and associated feeding behaviours. The key physiological pathways contributing to body weight gain and fat accumulation include neural mechanisms that drive food appetite and consumption, as well as the balance between fat utilisation and storage, collectively referred to as nutrient partitioning. Indeed, a diverse range of factors affect these energy-regulating neural mechanisms, including metabolic state (fasted vs fed), food availability, motivational state, emotional valence, reward processing and sensory awareness. Ultimately, the integration of these factors regulates food seeking, acquisition and nutrient partitioning. Therefore, novel approaches to treat or prevent obesity and other associated metabolic diseases need to incorporate an integrated understanding of the neural control of appetite and metabolism.

Traditionally, studies have focused on homeostatic interoceptive mechanisms, largely governed by the hypothalamus and brainstem. However, recent studies highlight that food consumption and nutrient partitioning are not solely dependent on the homeostatic requirements of the body. For example, the integration of external environmental cues, including the visual, auditory or olfactory perception of food availability and palatability is equally as important ^1, 2, 3^.

A variety of different neural structures process sensory information pertaining to food, which collectively help associate learned predictive cues with successful food acquisition and appropriate metabolic responses to match the predicted nutrient load. Importantly, homeostatic mechanisms can modulate the salience of sensory information depending on energy needs, suggesting an essential role for hunger-sensing and hunger signalling to integrate sensory information with energy need. For example, the sensory detection of food resets hypothalamic agouti-related peptide (AgRP) neurons and proopiomelanocortin (POMC) neurons within seconds, with a response magnitude that predicts the energy density of the food to be consumed and energy needed ^2, 4, 5, 6^. However, how neural hunger mechanisms influence sensory perception and detection of relevant environment cues remain unknown.

Although sensory information relating to food and food cues comes from a range of sources, olfaction is often the first sensory modality to assess food characteristics and prime behavioural and metabolic responses. In many animals, olfaction plays an essential role in food seeking ^7^ and food-related odours stimulate physiologic responses in anticipation of food (salivation, gastric acid secretion, lipid utilisation, blood pressure) ^1, 3, 8, 9^ to promote behavioural responses that increase appetitive motivation and memory formation ^7, 10^. Indeed, humans with olfactory dysfunction often report issues with appetite and enjoyment of food, highlighting an olfactory influence over the hedonic appreciation of food ^11, 12^. Notably, olfactory bulbectomy is a rodent model of depression-like behaviour ^13^.

The influence of olfaction on feeding behaviour and metabolism is exemplified by olfactory dysfunction in many metabolic diseases, including diabetes, obesity, and anorexia nervosa ^14, 15, 16^. Conversely, increased olfactory sensitivity prevented diet-induced obesity in both rodent genetic and pharmacological models ^17, 18, 19^. In addition, metabolic state regulates olfaction since hunger and energy deprivation increase olfactory discrimination and sensitivity ^20, 21, 22, 23^. Collectively, these studies point to an important, yet poorly understood, reciprocal relationship between olfaction and metabolism. One potential underpinning mechanism involves the action of circulating metabolic hormones that cross the blood-brain-barrier via fenestrated capillaries present in this vascularized region adjacent to the OB ^24^. Indeed, a vast array of hormone receptors are found in the OB including receptors for insulin, leptin, cholecystokinin, orexins, glucocorticoids and ghrelin ^15, 25, 26, 27^.

Given that hunger increases olfactory sensitivity ^20, 21, 22, 23^, we wanted to assess the mechanisms linking hunger with olfaction and the implications of disrupting this feedback on food intake, metabolism and related mood and foraging behaviours. A possible candidate linking hunger and olfactory function is ghrelin. Ghrelin is a metabolic hormone from the stomach conveying low body energy availability that helps maximize energy intake, energy storage, maintain blood glucose and modulate behaviours that facilitate energy-seeking during times of metabolic need ^28, 29^. In line with this, plasma ghrelin is increased during fasting, calorie restriction and starvation and reduced in diet-induced obesity ^30^, and diet-induced obesity is associated with a state of ghrelin resistance ^31, 32, 33, 34^. Ghrelin acts by binding to GHSRs (Growth Hormone Secretagogue Receptors) which are expressed throughout the CNS^35^, including the olfactory bulb and epithelium ^26, 27^. Evidence for OB GHSR expression includes, beta-galactosidase expression in the glomerular, mitral cell, and granular cell layers of the main OB and in the accessory OB of GHSR promoter-driven beta-galactosidase reporter mice ^27^, eGFP expression in the OB of GHSR-eGFP transgenic mice ^36, 37^ and GHSR mRNA expression in the OB of guinea pigs (as determined by RT-PCR) ^38^, as well as biotinylated ghrelin binding to scattered cells within the OB of sectioned rat brains ^27^. Indirect evidence for an important role of ghrelin to link hunger and olfaction comes from observations that the OB has either the highest or one of the top two highest uptakes of radiolabeled ghrelin following its i.v. administration in the entire brain ^39, 40^ and ghrelin influx across the blood-brain barrier is highest during fasting and lowest in obesity ^41^. Further, ghrelin administration markedly increases c-fos immunoreactivity within the mitral cell, inner plexiform, and granular cell layers of the OB ^42^, nasal application of ghrelin augments the percentage of c-fos-positive juxtaglomerular OB cells in mice induced by the odorant 2,3-hexanedione ^25^, and caloric restriction induces c-fos in new adult-born OB cells in a ghrelin-dependent manner ^36^, all of which suggest that ghrelin can activate OB cells. Although peripheral administration of ghrelin increases OB c-fos expression ^42^ or enhances food odour conditioning and sniffing in humans ^27^, the actions of endogenous ghrelin acting at its receptor in OB remain unknown. In this study, we hypothesized that GHSRs in the OB link hunger with olfactory sensitivity to control feeding-related behaviours and metabolism.

## Experimental Procedures

### Animals

All experiments were conducted in accordance with the Monash University Animal Ethics Committee guidelines and the Australian code for the care and use of animals for scientific purposes (8th edition) (2013). Male mice were kept under standard laboratory conditions with ad libitum access to food (20% protein 4.8% fat chow diet; Specialty Feeds, Western Australia) and water at 23°C with a relative humidity of 50-70% in a 12hr light/dark cycle. Mice were group housed (3-4 per cage) to prevent isolation stress unless otherwise stated. Mice were 10-12 weeks at time of surgery.

### Surgery

Floxed-GHSR (*Ghsr^fl/fl^*) mice were crossed with Ai14 mice (B6.Cg-*Gt(ROSA)26Sor^tm14(CAG-tdTomato)Hze^*/J; stock number #007914, Jackson Laboratory, Maine, USA) to develop a *Ghsr^fl/fl^*∷Ai14 RFP C57/BL6 mouse line. Ai14 mice are used to drive the expression of a red fluorescent protein (tdTomato) in the presence of cre. Targeted deletion of GHSR in the olfactory bulb (OB) was achieved through the bilateral stereotaxic injection of an Adeno-Associated virus containing a cre-recombinase enzyme (AAVpmSyn1-EBFP-Cre, Addgene#51507) into the OB of *Ghsr^fl/fl^*∷Ai14 RFP (designated OB^GHSR−/−^) and *Ghsr^wt/wt^*∷Ai14 RFP mice (designated WT) (coordinates: x= ± 0.5, y= 3.2, z= −1.5, −2.7). Mice were given two weeks to recover. Cre-driven RFP expression and Cre expression was confirmed via immunohistochemistry in all experimental mice post-mortem.

#### S*urgical procedure*

Mice were anaesthetised using isoflurane at a concentration of 5% for induction followed by 1-3% for maintenance. Anaesthesia was indicated by the loss of the pedal withdrawal reflex, after which mice were positioned in a stereotaxic apparatus. Subcutaneous injection of Metacam (50μl at 0.25mg/ml) was performed prior to surgery, and a drop of Lacri-Lube (Allergan) was applied to the eyes to prevent corneal damage during the surgery. An incision was made in the centre of the scalp to expose the bregma on the skull. Two 1-mm diameter holes were drilled in the skull with a drill at the bregma coordinates x= ± 0.5, y= 3.2. A 1μl Hamilton syringe was then lowered into the brain until reaching z=−2.7 to target the OB. An AAV-BFP-Cre virus (AAVpmSyn1-EBFP-Cre, Addgene#51507) was injected via the syringe into the tissue at a rate of 40nl/min for 5 min, and repeated at the z coordinate of −1.5, on both sides of the OB. Mice were given 30 minutes recovery time in a cage placed on a heat pad following surgery and monitored over a two-week period following the surgery. In that time, body weight, appearance, motility observation, and food intake were recorded.

### Olfactory Tests

#### Olfactory habituation test

This test is used to investigate olfactory detection and discrimination ability. Odours were presented in a sequential pattern with 3 trials per odour, 2 min exposure and 1 min apart. Odours were prepared in Eppendorf tubes with filter paper and five holes. A blank filter paper served as negative control, with a Froot loops (*Kellog’s cereal*) as a food-based odour with appetitive value, and rosewater as a novel non-food odour, allowing us to broadly assess the role of OB^GHSR^ deletion on olfactory performance. All mice were individually habituated to testing cages for 30 minutes prior to testing. Sniffing time, videoed and recorded by a stopwatch, indicated the level of interest in the odour and sniffing was only scored when the mice had their head oriented directly towards the Eppendorf tube and their nose was within 2 cm of the tube. Mice were tested under fed and 24-hour fasted conditions.

#### Buried Food Finding

This tests the olfactory performance of mice by measuring how much time is needed to find a buried familiar palatable food (latency); froot loop (*Kellog’s cereal*). All tests were performed four hours before the start of the dark phase in fed mice, 4-hour short fasted and 24-hour fasted conditions. The mice were given froot loops daily in their cages for three days prior to testing to acclimate them to the scent and minimise neophobia. Mice were placed in standard cages (27 x 18 cm) that mimicked testing conditions (4 cm layer of clean sawdust bedding) the night before each test to minimise stress and novel environment exploration. On the test day, mice were transferred to the testing room and transferred to an empty clean cage for 10 min to acclimatise to the room, while a froot loop was buried approximately 2 cm beneath the surface of the test cage containing the high bedding. The bedding surface was smoothed, and the mouse was then transferred to the test cage. The latency to find the froot loop (e.g. mouse eats or holds the froot loop) was videoed and recorded by an observer standing 2m away from the cage with a silent stopwatch. A cut-off of 6 min (fasted) or 10 min (fed) was used for the test.

#### Three Chamber Preference Test

This was used to assess olfactory preference. Mice were placed in the central compartment of the three-chamber apparatus containing two empty perforated Eppendorf tubes at opposite chambers and habituated for 10 min. Next, the two Eppendorf tubes were replaced with the other two Eppendorf tubes, containing filter paper, one scented with female urine, the other one a blank scent (6 min).

To assesses the preference for a social stimulus over a non-social stimulus, a similar three-chamber social interaction protocol was used. After the habituation phase with 2 empty wire mesh pencil cups, a social stimulus (female WT mouse) was introduced under one of the cups, while a novel object (scale weight) was introduced under the other cup (test phase 1, 6 min). In a second phase, the object stimulus was then replaced by a novel social stimulus (female WT mouse), and the test mouse was allowed to explore the arena for 6 min (test phase 2). The amount of time spent interacting with either stimulus was recorded.

### Behavioural Tests

All tests were performed ~4 hours before the start of the dark phase (14:00) under both fed and overnight fasted conditions. Behavioural experiments, excluding the saccharin preference test, were recorded, and analysed using Ethovision version 14.0.1322 (Noldus Information Technology; NL).

#### Open Field Test

This consisted of an open circular arena with a diameter of 80 cm. Within the open field, there was a centre zone, and a perimeter zone. Each mouse was initially placed in the same position of the perimeter zone and was then allowed to roam undisturbed in the arena for 5 minutes.

#### Light/Dark Box

This consisted of two different chambers: a large, illuminated ‘light box’ (48 x 30 cm), and a smaller, closed ‘dark box’ (15 × 30 cm). The chambers were separated by an open arch serving as a passageway. The mice were initially placed in the dark box and were then allowed to roam undisturbed in the apparatus for 5 minutes.

#### Elevated Plus Maze (EPM)

The EPM consisted of two open arms and two closed arms (5 × 30 cm) that intersected at 90 degrees to form a plus sign with a centre zone (5 x 5 cm) in the middle. It was raised 50cm from the ground. The mice were initially placed in the central zone facing the north open arm and were then allowed to roam undisturbed through the maze for 5 minutes.

#### Saccharin Preference Test

This was used to test sensitivity to reward, in which singly housed mice were offered either water or a palatable non caloric solution of 0.1% Saccharin for 2 hours every day (14.00 to 16.00) for 4 days. Preference score was calculated by the saccharin intake (V_s_) normalized to water intake (V_w_): Preference score (%) = V_s_/(V_s_ + V_w_)*100

#### Spontaneous alternation test

This test is driven by innate curiosity to explore novel areas and measures locomotor and exploration activity, as well as spatial working memory. Mice were not habituated but placed in the room before the test. A novel Y-maze was used to drive spontaneous exploration. Mice were allowed to explore 3 arms of the Y maze during which the sequences of entries are recorded. Alternation is determined from successive entries of the 3 arms on overlapping triplet sets in which 3 different arms are entered, for example the arm sequence ACBABACBAB results in 5 alternations: ACB, CBA, BAC, ACB, CBA. The number of alternations is then divided by the number of alternation opportunities, namely total arm entries minus one (Spontaneous alternation behaviour score, SAB score).

#### Y Maze Exploration

The Y-shaped maze was set-up in the room with visual cues on walls. In the first trial the mice were able to explore the maze for 10 min while 1 arm was blocked. In the second trial all arms of the maze were open, and mice are allowed to explore all arms of the maze for 5 min. The Inter-trial interval was 30 min. The entry into the previously blocked arm indicates the willingness to explore the new arm.

#### Baited Y Maze

This tests cognitive function based on the ability of the mouse to remember spatial locations. The Y-shaped maze was set-up in the room with visual cues on walls, as well as intra-maze cues. Mice were given peanut butter chips in their home cages prior to testing to prevent food neophobia and were habituated with the Y-maze the day before to become familiar with the maze as well as to measure initial preference. Following 2x 2-minute training sessions, where a peanut butter chip (*Reese’s*) is placed in one of the 2 test arms, mice were allowed to explore the two arms of the Y-maze (test) and exploration of the previously baited arm was measured. The test was performed 2 hours following the final training session.

#### Radial Arm Maze (RAM)

RAM measures spatial learning and memory. The apparatus consisted of 8 arms with visual cues, inside and outside of the maze. Mice were habituated the day before for 10 min with no reward in the maze. On the test day working memory was assessed. A reward (peanut butter chip, 50-70 mg) was placed into each arm of the RAM. Working memory is then assessed by examining entries into each arm with re-entries into a previously entered arm resulting in a working memory error. The memory score is calculated by the difference of the correct arm entries to incorrect entries divided by the total arm entries.

#### Delayed spatial win-shift test in RAM apparatus

In this test the rewarding arms during the trial and test phase is alternated or shifted, while no more than two adjacent arms can be blocked or baited. The mouse is required to hold spatial information both within the task and across the delay (inter-trial interval with mouse in home-cage: 2 min) to obtain rewards (peanut butter drop) and measures the type and incidence of memory errors made in the training (4 min) and test phase (4 min) of the learning task. A working memory error (re-entry of an arm that has been baited) can occur in both phases of the task, while a reference memory error (entry into an arm that has been baited during the training phase and is no longer baited) can only occur during the test phase. Mice conducted a total of 14 trials within 3 weeks, and consecutive trials within one week were binned to observe their learning progress. The night before the test, access to food was restricted to 80% of normal food intake to encourage food-seeking behaviour in the maze. The memory score is calculated by the difference of the correct arm entries to incorrect entries divided by the total arm entries.

### Food Intake Experiments

Food intake was measured in single-housed mice after a short (4 hour) or 24 hour fast and chow food intake was measured at 1, 2, and 5-hour timepoints after approximately 20g of chow was reintroduced to the cages manually. For the intraperitoneal (IP) ghrelin-induced feeding experiment, mice were given either a ghrelin injection (0.5 μg per g bodyweight) or a saline (0.9%) injection of the same volume. In addition, mice were housed in an automated feeding monitoring system (BioDaq Feeding Cages, Research Diets, NJ, USA) to measure episodic ad-libitum feeding activity and behaviour of singly housed undisturbed mice.

### Glucose tolerance test (GTT) and Insulin tolerance test (ITT), 2-Deoxyglucose Challenge

The mice were fasted 4 hours before testing in the light phase at 13:00. Fasting blood glucose (t= 0 min) was measured using an ACCU-CHEK blood glucose monitor. For the GTT, glucose was diluted in tap water to a concentration of 0.25 g/ml, and was delivered via oral gavage at a dose of 2mg/g bodyweight. The IP insulin dose during the ITT delivered was 0.75 mU/g bodyweight. 2-Deoxyglucose (2-DG, *Sigma #D6134*) was administered via IP injection at 0.5 mg/g bodyweight. Blood glucose was measured at 15, 30, 60, 120 min. Blood samples (approx. 20μl) were collected in tubes containing EDTA at t= 0 min, t= 30 min, and t= 60 min. The course of insulin secretion during the GTT was determined. The counterregulatory response to insulin or 2-DG-induced hypoglycaemia was assessed by measuring corticosterone secretion.

### 24-Hour Fasting Blood Glucose

After fed blood glucose at t= 0 (09:00) was measured, food was removed from the cages and blood glucose was measured over a 24-hour window as indicated, and then the following day at 09:00. Blood samples (approx. 20μl) were collected in tubes containing EDTA.

### Gastric emptying during the GTT

Fasted mice (4h) were administered glucose and acetaminophen via an oral gavage dose at 2g/kg glucose and 0.1g/kg acetaminophen. Blood glucose and acetaminophen were measured over a 2h time period. Since acetaminophen only enters the blood stream after it passes the stomach, appearance in blood can be used as an estimate for gastric emptying. Plasma acetaminophen was measured via colorimetric detection kit (2k99-20, Abbott).

### Faeces Triglycerides

Faecal lipids were extracted using the principle of a Folch Extraction protocol. In brief, faecal samples of a 24h period were collected and total faecal weight per day recorded. 100 mg of faeces was used for analysis, first softened in 400μl ddH2O, then 1.5ml of cold chloroform:methanol (2:1) mixture added and lysed using a Qiagen bead TissueLyser, mixed at RT for 20 min prior to centrifugation for 30 min at 9000 rpm. The lower liquid phase was transferred to a fresh tube containing 200μl 0.9% NaCl, centrifuged for 5 min at 2000 rpm. The lower organic phase was transferred into 40μl of chloroform:Triton-X (1:1) solution, dried with a speedvac overnight. To measure lipids, 200μl of ddH2O was added to the remaining triton-lipid solution and a TG (Sigma) or NEFA assay kit (Novachem) was used.

### Indirect Calorimetry

Metabolic parameters, oxygen consumption (VO2; ml/hr), carbon dioxide production (VCO2; ml/hr), respiratory exchange ratio (RER), energy expenditure (EE; kcal/hr), and locomotor activity were measured using the Promethion system (Sable Systems International). Mice were single-housed and had *ad libitum* access to food and water in overhead feeders attached to electronic balances to measure food and water intake. No data were acquired in the respiratory cages for 5 days to permit acclimation to their new environment. Baseline metabolism was assessed within a 48-hour period.

### Hormone Level Analyses

Plasma insulin concentrations were determined using an ELISA (CrystalChem #90080) according to manufacturer’s instructions. Corticosterone concentrations were determined using an ELISA (Abcam #ab108821) according to manufacturer’s instructions.

### PCR and primers

Total RNA was extracted using a guanidium-phenol-chloroform phase separation method, according to the Qiazol® Lysis Reagent (Qiagen #79306) manufacturer’s protocol, followed by precipitation of the RNA pellet using isopropanol, 10 min incubation at room temperature and pelleted at 12,000×g, 4°C, 10 min. The pellet was washed 3 times with 75% ethanol prior to reconstitution in RNAfree water. The relative purity and concentration of the RNA was determined using a QIAEXPERT spectrophotometer. Total RNA was treated with DNase I (Qiagen #79254), and complementary DNA (cDNA) was synthesized from RNA using iScript cDNA Synthesis Kit (Bio-Rad #1708891). For qPCR, Fast SybrGreen (Thermofisher Scientific #4385617) was used for amplification and detection, with 25 ng cDNA loaded per well. The Rotor-Gene Q real-time PCR cycler (Qiagen) was used to determine the mRNA expression level for the genes of interest. Following validated primers were used: GHSR: Forward 5’-GCTGCTCACCGTGATGGTAT-3’, Reverse 5’-ACCACAGCAAGCATCTTCACT-3’, Actin: Forward 5’-CCAGATCATGTTTGAGACCTTC-3’, Reverse: 5’-CATGAGGTAGTCTGTCAGGTCC-3’

### Western Blot

Proteins were detected by Western blot analysis after separating 30–50 μg of total protein lysate prepared in RIPA buffer with protease inhibitor cocktail (cOmplete™, Roche) on a 4– 15% Mini-PROTEAN® TGX™ Precast Gel (BioRad) and transferring to a polyvinylidene difluoride membrane (Immobilon-P 0.45 μm, Millipore). The blots were blocked for 1 hour with 3% BSA and then probed with GHSR Polyclonal Antibody (ThermoFisher # PA5-28752,1:1000 dilution) or anti-Actin (Sigma #A2066) after stripping of the membrane and detected via enhanced chemiluminescence (ECL, Sigma). Westerns were analysed using ImageLab Software (BioRad). A reference sample was including in all blots and GHSR protein expression was normalized to actin.

### Immunohistochemistry

To confirm viral injection, animals were deeply anesthetized with isoflurane and perfused with 0.05 M PBS, followed by 4% paraformaldehyde. Brains were postfixed in 4% paraformaldehyde overnight at 4°C, then placed in 30% sucrose. Brains were cut at 40 μm on a cryostat, and every fourth section was collected and stored in cryoprotectant at −20°C. Sections were washed in 0.1 M phosphate buffer (PB), incubated with 1% hydrogen peroxide (H2O2) for 15 minutes to quench endogenous peroxidase activity, and blocked for 1 hour with 5% normal horse serum (NHS) in 0.3% Triton in 0.1 M PB. Sections were incubated with rabbit anti-DsRed (TaKaRa Bio, cat. no. 632496) at 1:1000 and guinea pig anti-Cre (Synaptic Systems, cat. No. 257 004) in diluent of 5% NHS in 0.3% Triton in 0.1 M PB. After incubation, the sections were washed and incubated with Alexa Fluor 594 donkey anti-rabbit antibody and Alexa Fluor 488 anti-guinea pig (ThermoFisher) at 1:2000 in 0.3% Triton in 0.1 M PB. Sections were then washed, mounted, and coverslipped.

### Statistical analysis

Statistical analyses were performed using GraphPad Prism for MacOS X. Data are represented as mean ± SEM. Two-way ANOVAs with post hoc tests were used to determine statistical significance. A two-tailed Student’s paired or unpaired t-test (see figure legends for specific details) was used when comparing genotype only. p < 0.05 was considered statistically significant and is indicated on figures and in figure legends.

## Results

### Deletion of OB^GHSR^

We first determined the effects of a 14-hr overnight fast and 2 hrs of re-feeding on GHSR mRNA expression in the OB (OB^GHSR^) and hypothalamus of WT mice. Fasted mice exhibited a 7.5-fold and 2-fold increases in GHSR expression in the OB and hypothalamus, respectively, over that observed in ad lib-fed mice; re-feeding normalized the levels **(Fig. 1A)**. Thus, hunger regulates OB^GHSR^ gene expression, highlighting it may be an important target to manipulate hunger signalling in the OB. To assess the role of OB^GHSR^ in feeding-related behaviours and metabolism, we generated a temporal and site specific deletion of GHSRs in the OB (OB^GHSR−/−^) of adult mice using bilateral stereotaxic injections of AAV-Cre into *Ghsr^fl/fl^*∷Ai14 RFP (OB^GHSR−/−^) and *Ghsr^wt/wt^*∷Ai14 RFP mice (WT) mice (**Fig. 1 C**). Cre recombinase-induced expression of red fluorescence protein (RFP) was observed in main olfactory bulb in both the granular and mitral cell layer, anterior olfactory nucleus and accessory olfactory bulb ranging from bregma 4.28 to 2.68 (**Fig. 1C**, **Suppl. Figure 1**), in both OB^GHSR−/−^ and WT respectively, confirming accuracy of injections to broadly cover the OB (**Fig. 1C**). Moreover, GHSR protein expression in the OB (from bregma 4.0 to bregma 2.6) was significantly reduced in OB^GHSR−/−^ compared to WT controls (**Fig. 1B**).

**Figure 1.**
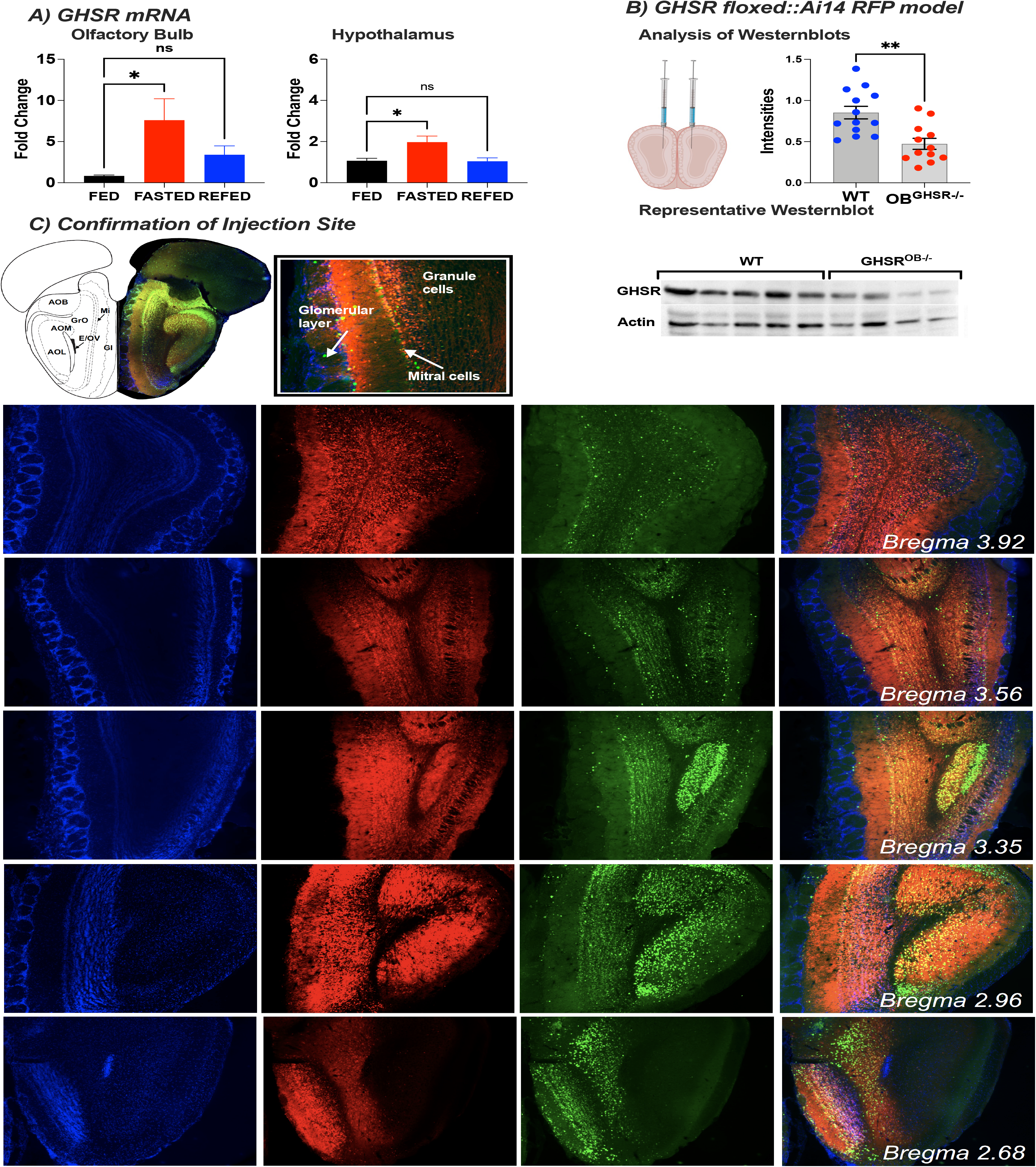
GHSR expression in the olfactory bulb during different metabolic conditions and after deletion. **A)** GHSR mRNA expression in the olfactory bulb and hypothalamus changes during fed (n=7/7), fasted (n=11/15) and refed (n=5/6) conditions. **B)** Confirmation of GHSR deletion in the olfactory bulb. Deletion efficacy after viral Cre recombination in the GHSR floxed∷Ai14 RFP model was confirmed by Western blot analysis. Protein GHSR expression intensities of 3 blots (top, WT=13, OB^GHSR−/−^=12) were normalized to corresponding actin. Representative blot (below). **C)** Immunohistochemistry for confirmation of injection site. Cre recombinase induced expression of red fluorescence protein (RFP, red) and Cre (green) in the olfactory bulb ranging from bregma 4.28 to 2.68, in both OB^GHSR−/−^ and WT respectively.

### OB^GHSR^ affects olfaction

To assess whether OB^GHSR^ deletion affects olfactory function, we used various olfactory performance tests in fed and overnight fasted conditions. In an olfactory habituation test (**Fig. 2A**), OB^GHSR^ deletion significantly reduced sniffing time to froot loops and rosewater and although fasting increased sniffing time compared to fed mice in both groups, this was still significantly attenuated in OB^GHSR−/−^ mice (**Fig. 2B-F**). Video analysis of mice behaviour during the olfactory habituation test revealed that OB^GHSR−/−^ mice spent less time in active states, such as walking, climbing, rearing and more time in stationary behaviour over the entire testing period (all trials; **Fig. 2G-J**). Furthermore, OB^GHSR−/−^ mice exhibited fewer behavioural changes during the olfactory habituation test (**Suppl Fig. 2**), suggesting less engagement and behavioural flexibility in exploratory activities. This was similar in an odour-baited field test, in which OB^GHSR−/−^ mice spent less time sniffing peanut butter scented paper in an open field environment (**Fig. 3A-G**). Although time spent in the inner zone was significantly reduced in both fed and fasted OB^GHSR−/−^ mice, only fasted OB^GHSR−/−^ mice spent less time in the sniffing zone. Behavioural analysis during the baited open field test revealed that OB^GHSR−/−^ mice show different behavioural patterns with more stationary behaviour and less involvement in sniffing and walking exploration (**Fig. 3F-G**) or behavioural transitions (**Suppl Fig. 3A-F**).

**Figure 2.**
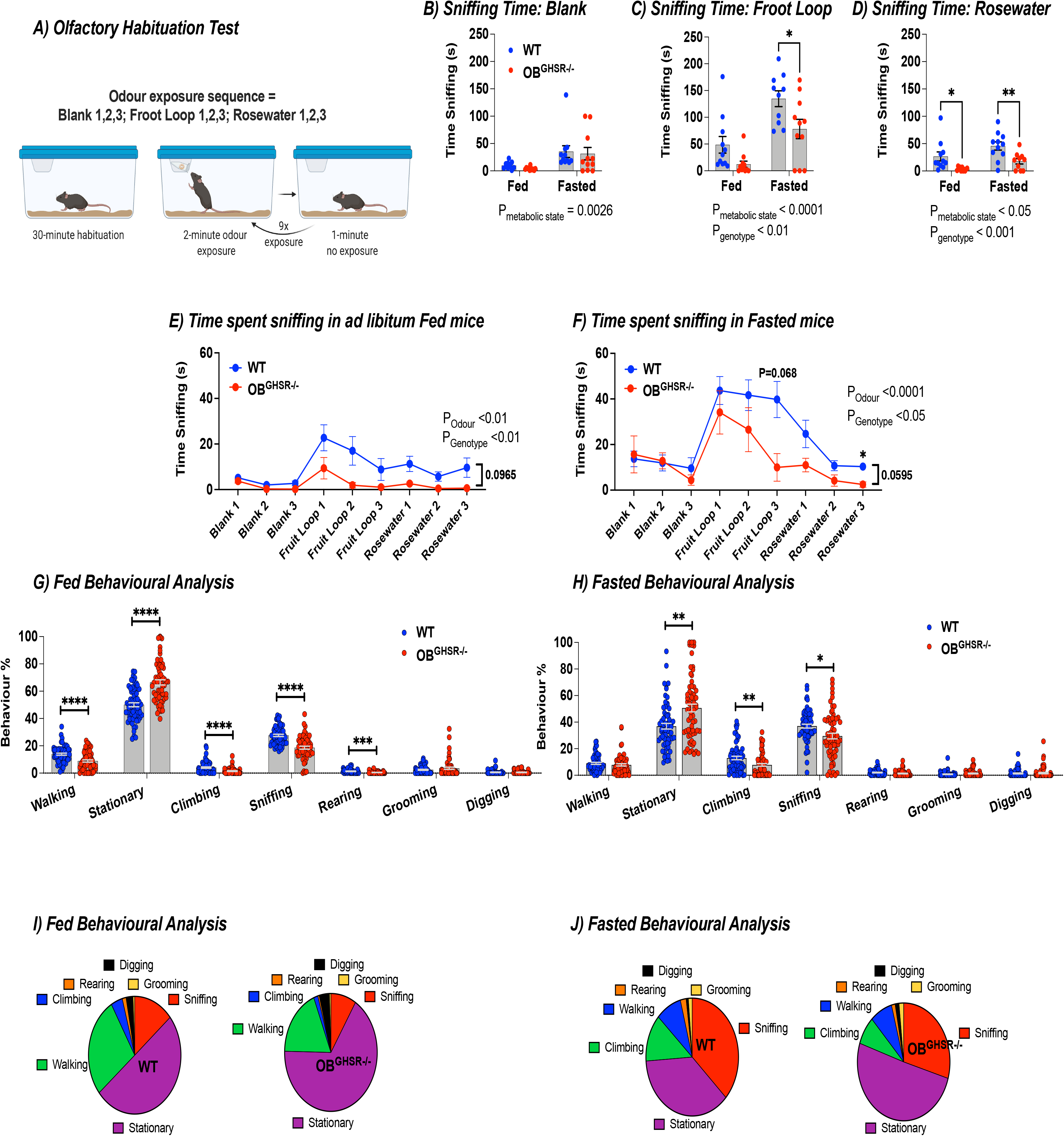
Reduced olfactory performance after OB-GHSR deletion in olfactory habituation test. **A)** Schematic of the olfactory habituation task created with BioRender.com. Mice were acclimatised to testing conditions for 30 min, then exposed to an odour for 3 consecutive times for 2 min each separated by 1 min (WT=11, OB^GHSR−/−^=11). Odour was presented on filter paper inside an Eppendorf tube with a perforated lid. **B-D)** Sniffing time fed versus fasted (B) Blank (Two-way ANOVA, P_metabolic state_=0.0026, P_genotype_=0.5631), (C) Froot Loop (Two-way ANOVA, P_metabolic state_<0.0001, P_genotype_=0.0024), (D) Rosewater (Two-way ANOVA, P_metabolic state_=0.0145, P_genotype_=0.0005). **E-F)** Time spent sniffing in (E) ad libitum fed condition across the 3 different odour exposures (Two-way ANOVA, P_odour_=0.0026, P_genotype_=0.0036), versus (F) fasted condition (Two-way ANOVA, P_odour_<0.0001, P_genotype_=0.0482, P_odour*genotype_=0.0595). **G-H)** Behavioural analysis during the olfactory habituation test in (G) fed, versus (H) fasted condition, combined for foot loop and rosewater trials. Behavioural differences were observed in exploratory behaviours, such as walking, climbing, sniffing, rearing, and stationary behaviours in the (G) fed state (P=0.000097, P=0.000121, P<0.000001, P=0.000304, P<0.000001), as well as for climbing, sniffing and stationary behaviour in the (H) fasted state (P=0.009947, P=0.021389, P=0.001596). **I-J)** Pie charts of behaviours (walking, climbing, rearing, digging, grooming, sniffing, stationary) during (I) fed, and (J) fasted conditions.

**Figure 3.**
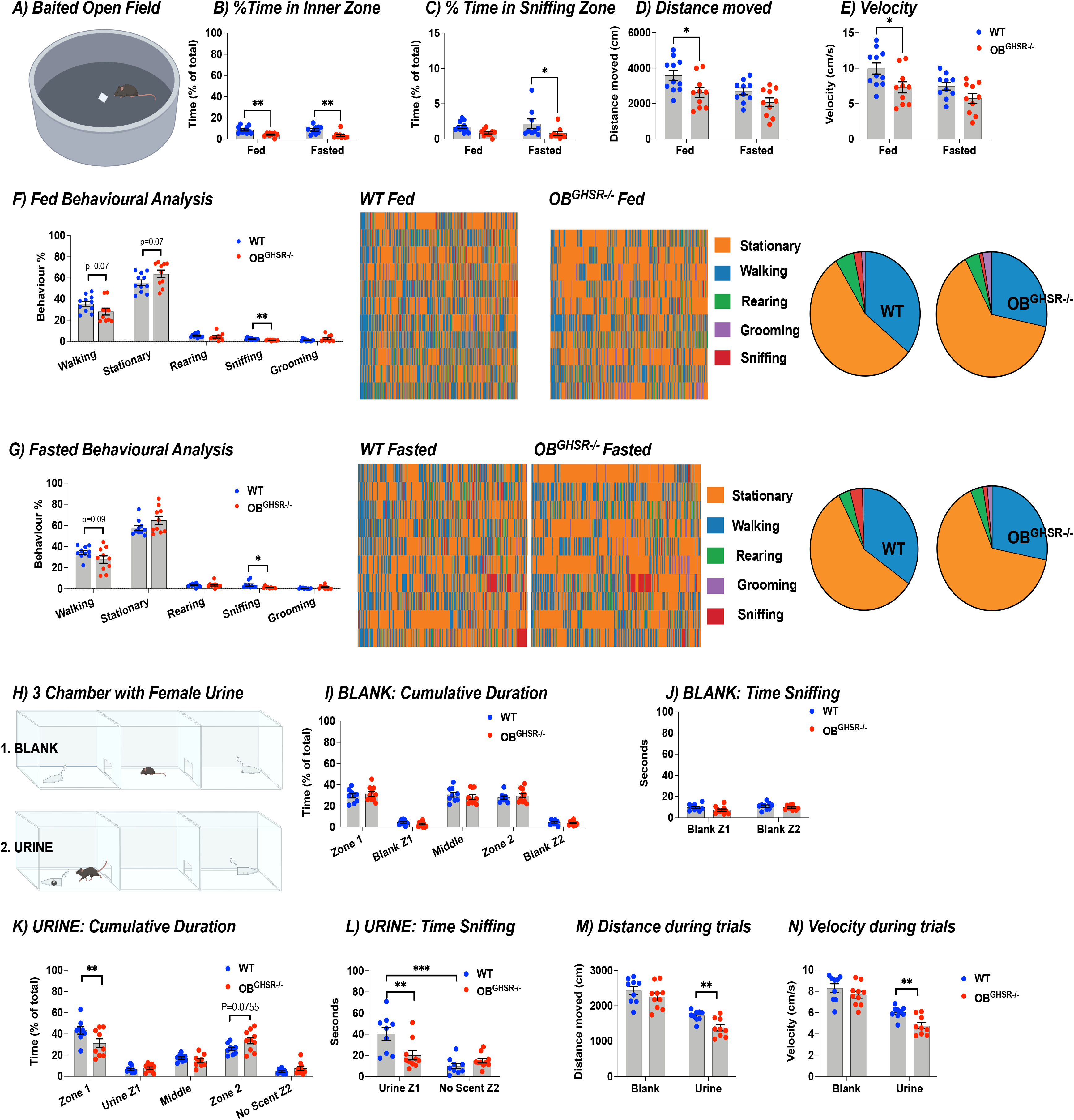
OB-GHSR deletion causes reduced interest in food-related odour as well as pheromones. **A)** Schematic of the baited open field test created with BioRender.com. A peanut butter-scented filter paper was placed in the middle of an open field arena, and mice were allowed to roam undisturbed in fed versus fasted conditions (WT=11, OB^GHSR−/−^=10). **B-E)** Percent Time in Inner Zone (B), and in sniffing zone (C), with the distance moved (D), as well as velocity (E). OB^GHSR−/−^ mice spend less time in the inner zone (Two-way ANOVA, P_genotype_<0.0001), and move less (Two-way ANOVA, P_genotype_=0.0039). **F,G)** Behavioural analysis with pie charts of behaviours (stationary, walking, rearing, grooming, sniffing) in fed (F) and fasted (G) conditions. Behavioural analysis during the baited open field task reveals that OB^GHSR−/−^ mice have less interest in active behaviours and spent more time stationary (multiple unpaired t tests, *Suppl. Table 1*). **H)** Schematic of the 3 chamber exploration with no scent or female urine on filter paper inside an Eppendorf tube with a perforated lid created with BioRender.com (WT=9, OB^GHSR−/−^=10). **I,J**) Cumulative duration (I) and time sniffing (J) in the different zones with no scent (blank) present. **K,L)** Cumulative duration (K) and time sniffing (J) in the different zones when mice were presented with female urine. **M)** Distance during trials. **N)** Velocity during trials. While there was no difference in the exploration task when no scent was present, OB^GHSR−/−^ mice spent significantly less time sniffing the female urine (Two-way ANOVA, P_genotype*zone_=0.0008) and explore less (multiple unpaired t tests, P_urine_ =0.002697, *Suppl. Table 1*).

Since olfaction is important for social behaviour, we used a three-chamber preference protocol to assess interest in non food-related social olfactory stimuli (**Fig. 3H-N**). Although time spent sniffing blank tubes in the habituation phase did not differ between control and experimental groups (**Fig. 3I-J**), OB^GHSR−/−^ mice spent less time in the chamber with the female mouse urine scent and sniffed less (**Fig. 3K-L**). Both distance moved and velocity were significantly lower in OB^GHSR−/−^ mice in the chamber paired with urine scent (**Fig. 3M-N**). A similar protocol was used to investigate social interaction by introducing a female mouse under and mesh cup versus a novel object (test phase 1) or subsequently a novel female mouse (test phase 2). No difference between the groups were observed (**Suppl Fig. 3G-M**), suggesting the additional visual cues made up for the lack of olfactory deficits. Collectively, these experiments demonstrate OB^GHSR^ deletion has a profound effect to decrease olfactory habituation, preference and sensitivity to both food and non-food odours, particularly in the fasted state.

### OB^GHSR^ regulate exploratory activity and mood

In the olfactory experiments described above we observed numerous indicators of impaired exploratory behaviour OB^GHSR−/−^ mice, suggesting potentially increased anxiety-like behaviour. To further investigate anxiety-like behaviour in OB^GHSR−/−^ mice, we assess behaviour using elevated plus maze (**Fig. 4A**), a light-dark box (**Fig. 4E**), and the open field (**Fig. 4I**) tests. In the elevated plus maze OB^GHSR−/−^ mice spent more time in the closed arms and moved less in the fasted state (**Fig. 4B-D**). This was similar for the light-dark box, where OB^GHSR−/−^ mice spent less time in the light zone with fewer light zone entries and moved less in the fasted state (**Fig. 4F-H**). Similarly, in the open field arena, fasted OB^GHSR−/−^ mice spent less time in the inner zone, with fewer inner zone entries and moved less (**Fig. 4J-L, Suppl. Fig. 4**). Of note, the differences in anxiety-like behaviours were not due to a generalized impairment in locomotor activity measured using home-cage running wheels (**Fig. 4M-O**). Anxiety-like behaviour is often linked with depression-like behaviours in mice ^43, 44^ and OB bulbectomy is a rodent model of depression ^13^, therefore we used a saccharin preference test of anhedonia to assess depression-like behaviour. Anhedonia is the inability to experience pleasure and we observed that OB^GHSR−/−^ mice consumed less saccharin over 4 days (**Fig. 4P-R**). Taken together, these results suggest that the OB^GHSR^ deletion results in greater depression-like and anxiety-like behaviour, the latter of which is more pronounced in the fasted state.

**Figure 4.**
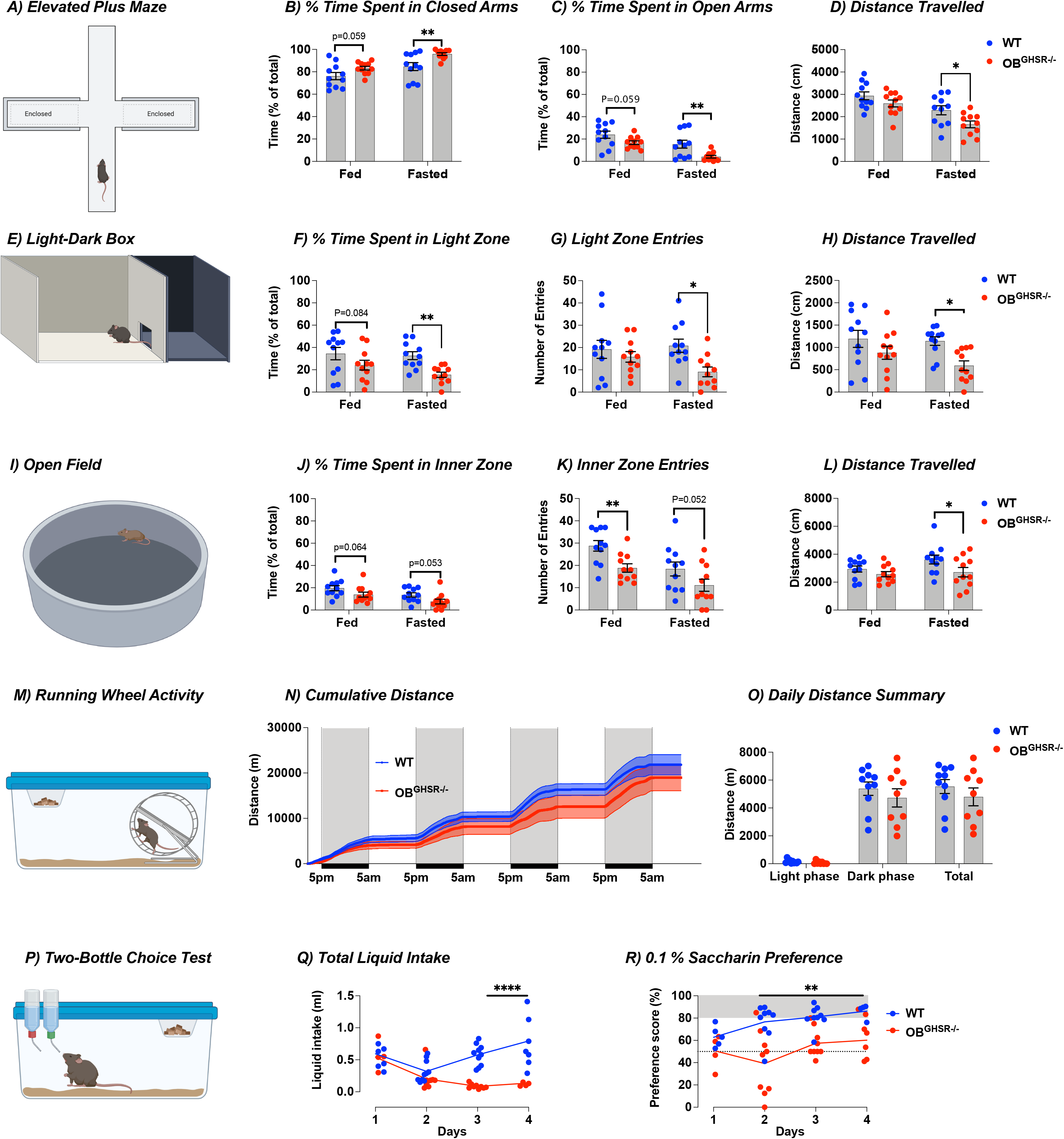
OB GHSR deletion affects mood, anxiety, and hedonia. **A-D)** Schematic of the elevated plus maze with 2 open arms and 2 closed arms in fed and fasted conditions created with BioRender.com (WT=11, OB^GHSR−/−^=11). Time spent in the more anxiogenic open zone was measured in fed versus fasted conditions. While mice with GHSR deletion spent more time in closed arms (B) and less time in open arms (C), this was significant in the fasted state, where mice also moved less (D) (Two-way ANOVA, *Suppl. Table 1*). **E-H)** Schematic of the Light-Dark Box in fed and fasted conditions created with BioRender.com (WT=11, OB^GHSR−/−^=11). Percent time in the anxiogenic light zone (F) and light zone entries (G) were significantly lower in OB^GHSR−/−^ mice when fasted, also with less distance travelled (H) during fasted conditions (Two-way ANOVA, *Suppl. Table 1*). **I-L)** Schematic of the Open Field Test in fed and fasted conditions created with BioRender.com (WT=11, OB^GHSR−/−^=11). Percent time (J) in the anxiogenic inner zone tended to be decreased in both metabolic states with fewer inner zone entries (K), and less distance travelled (L) (Two-way ANOVA, *Suppl. Table 1*). **M-O**) Schematic of the home cage running wheel activity created with BioRender.com. Cumulative distance (Q) and daily distance summary (R) for 4 days shows no difference between the genotypes (multiple unpaired t tests, *Suppl. Table 1*). **P-R)** Schematic of the Two-Bottle Choice Test created with BioRender.com. Mice were offered either water or a palatable non-caloric 0.1% Saccharin solution for 2 hours every day (WT=10, OB^GHSR−/−^=9). OB^GHSR−/−^ mice have a significantly lower saccharin intake (N) and calculated saccharin preference score (O) (Two-way ANOVA, *Suppl. Table 1*).

### OB^GHSR^ regulate food-seeking but not food intake

Since ghrelin administration increases food intake, we performed feeding experiments to investigate the effects of OB^GHSR^ deletion on food seeking and consumption. Although total cumulative food intake was not significantly different (**Fig. 5A-C, Suppl Fig. 5A**), OB^GHSR−/−^ mice displayed significant differences in feeding behaviour with fewer feeding bouts during the dark phase and after fasting (**Fig. 5D-F, Suppl. Fig. 5B,D**), and spent more time per bout (**Fig. 5G**). Moreover, food intake was not different after fasting or after ghrelin injection (**Fig. 5H,I, Suppl Fig. 5C**). However, in a buried food-seeking test often used to assess olfactory capacity (Machado, 2018 #151), OB^GHSR−/−^ mice took significantly longer to find a froot loop in response to a short 4-hour fast (**Fig. 5K**) or IP ghrelin compared to WT (**Fig. 5M**), and while this difference was not present in the fed state (**Suppl. Fig. 5E**) a strong trend was also observed after overnight fasting (p=0.079; **Suppl Fig. 5F**). Collectively, these results suggest OB^GHSR^ does not affect daily ab libitum food intake, or in response to fasting and IP ghrelin, but rather significantly impairs normal feeding behaviour and food seeking.

**Figure 5.**
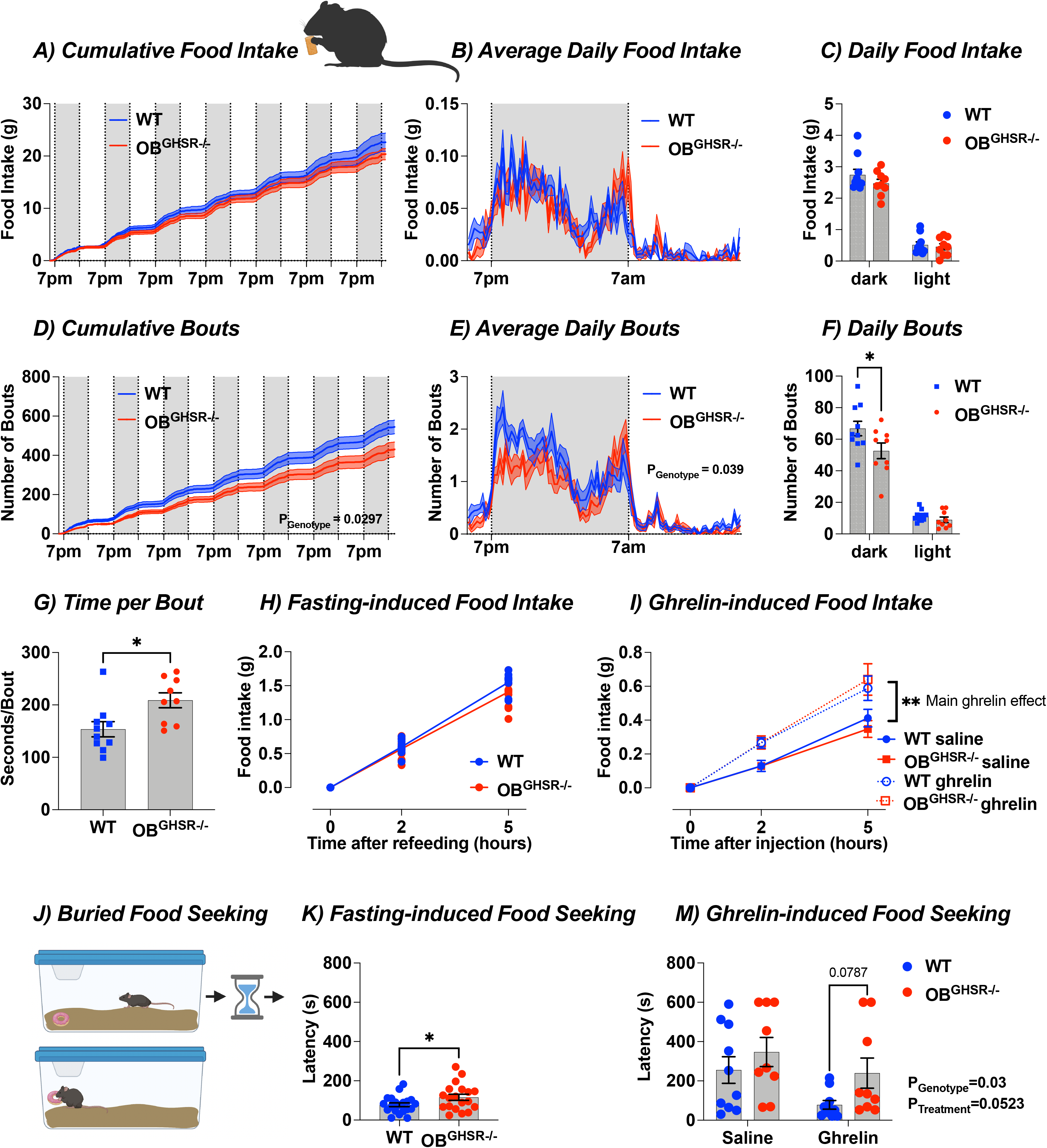
OB GHSR deletion does not affect food intake but feeding behaviour and food seeking. **A-G)** Food intake in BioDaq Feeding Cages (WT=11, OB^GHSR−/−^=11). Cumulative Food Intake (A), with average daily food intake (B), and daily food intake during the dark and light phase (C) during a 7-day period. Average food intake was not different between the genotypes (Two-way ANOVA, *Suppl. Table 1*). However, ingestive behaviour was different between the groups as shown with cumulative bouts (D), average daily bouts (E), daily bouts (F), and time spent per bout (G). OB^GHSR−/−^ mice had significantly fewer feeding bouts (Two-way ANOVA, *Suppl. Table* 1) and spent more time per bout (Two-tailed unpaired t test, P=0.0155). **H)** Fasting-induced food intake (WT=11, OB^GHSR−/−^=11). Food intake in WT and OB^GHSR−/−^ following an overnight fast 2h and 5h after the food was reintroduced. There was no significant different between the groups (Two-way ANOVA, *Suppl. Table 1*). **I)** Ghrelin-induced food intake (WT_saline_=18, WT_ghrelin_=20, OB^GHSR−/−^ _saline_=21,OB^GHSR−/−^ _ghrelin_ =20). Food intake in WT and OB^GHSR−/−^ following intraperitoneal injection of ghrelin (0.5 μg/g). While we observed a significant effect of ghrelin on food intake, there was no difference between the two genotypes (Two-way ANOVA, *Suppl. Table 1*). **J-M)** Schematic of the Buried Food Seeking Test created with BioRender.com, which records the time needed to find familiar palatable food (froot loop) buried in high bedding. **K)** Fasting-induce food-seeking after a 4-hour fast (WT=20, OB^GHSR−/−^=19). OB^GHSR−/−^ mice took significantly longer time to find the froot loop (Two-tailed unpaired t test, P=0.0428). **M)** Ghrelin-induced food seeking (WT_saline_=10, WT_ghrelin_=10, OB^GHSR−/−^_saline_=9, OBGHSR−/−_ghrelin_ =9). OB^GHSR−/−^ mice needed more time to find the froot loop (Two-way ANOVA, P_genotype_=0.03, *Suppl. Table 1*).

### OB^GHSR^ deletion does not affect spatial memory and learning

Olfactory cues are often used in spatial navigation and food or food cues are often used to guide spatial exploration tasks ^45, 46^. To assess whether OB^GHSR^ contribute to spatial exploration and learning, we used a Y maze and radial arm maze (RAM). In general, OB^GHSR−/−^ mice moved less during exploration tasks in a Y maze, as reflected by less distance moved in the sequential alternation test (Fig. **6A,D**) with few entries into the triangular transition zone and arms (**Fig. 6B,C**) and thus fewer alternations (the sequential entry of 3 different arms) (**Fig. 6F**). However, when looking at the number of alternations compared to the total number of alternation opportunities, there was no difference in the SAB score (**Fig. 6G**), with no difference in inter-choice interval (**Fig. 6E**), suggesting deletion of GHSR in the OB did not alter short term spatial memory. In a novel spatial exploration Y-maze task (**Fig. 6H**), although OB^GHSR−/−^ mice had fewer novel arm entries and spent less time in the novel arm, they also moved less in the Y maze (**Fig. 6I-K**), similar to anxiety-like behavioural tests (see Fig. 4D,H,L). Therefore, to incentivise movement and spatial exploration we used a baited Y maze approach during the fasted state, where mice had to memorize the reward location in the Y maze during two training sessions (**Fig. 6L**). Moreover, mice were habituated to the Y maze to minimise any potential anxiety-like behavioural impairments. Under test conditions and in the absence of food, OB^GHSR−/−^ mice entered the previously baited arm with similar latency, spent similar amount of time in that arm, and exhibited similar preference scores as controls in fasted (**Fig. 6M-O**) and ad libitum fed conditions (**Suppl. Fig. 6B,C,E**).

**Figure 6.**
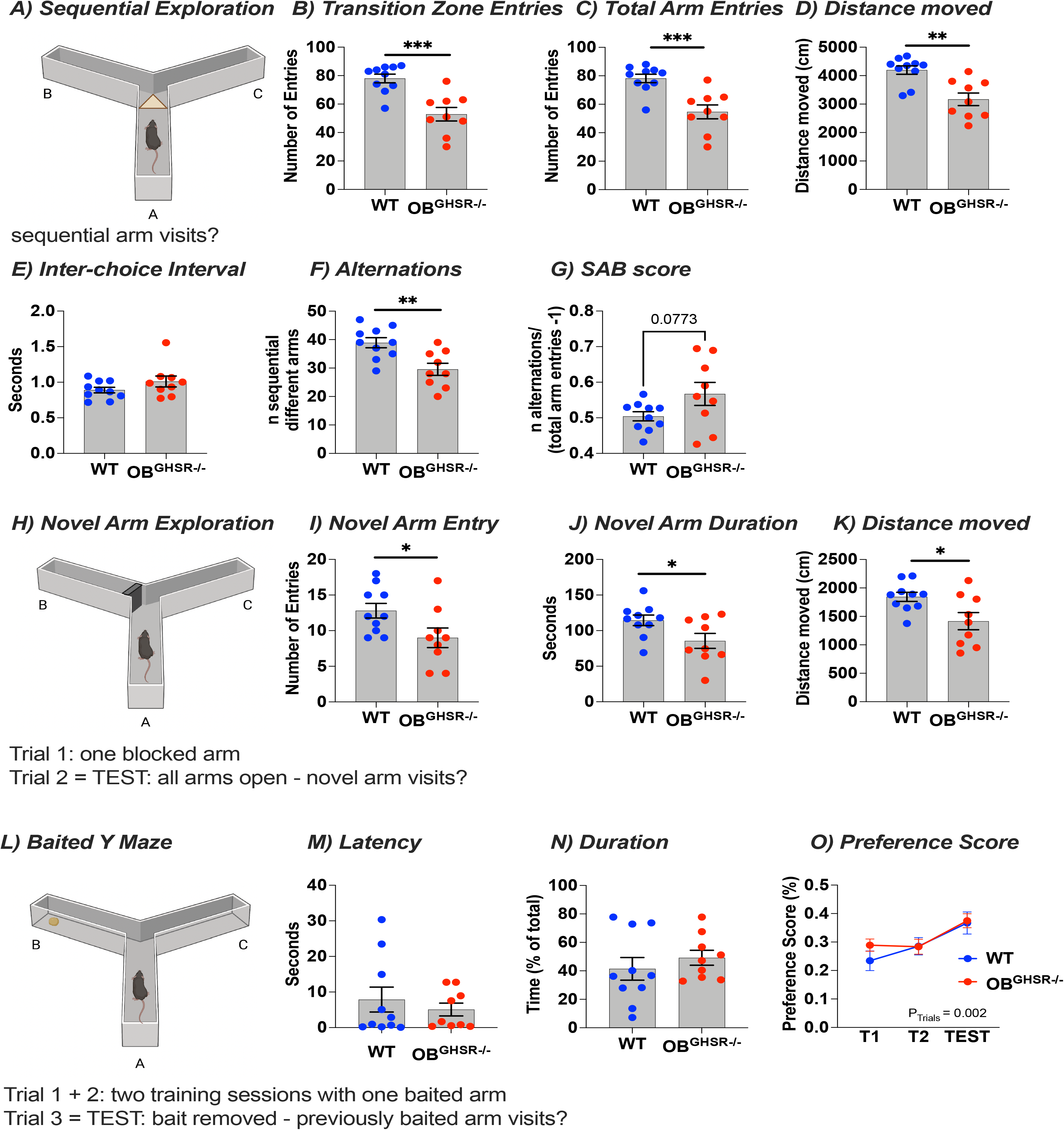
OB GHSR deletion does not affect spatial exploration. **A-G)** A schematic of the Sequential alternation test created with BioRender.com, which measures the spontaneous exploration activity of a mouse in the Y-maze (WT=10, OB^GHSR−/−^=9). OB^GHSR−/−^ mice moved less during exploration tasks in a Y maze, as reflected by fewer transition zone entries (B), which is the triangle in the middle of overlapping arms; fewer total arm entries (C), and total distance moved (D) (Two-tailed unpaired t tests, *Suppl. Table 1*). **E)** Inter-choice interval or time spent in the transition zone (Two-tailed unpaired t tests, *Suppl. Table 1*). **F)** Alternations: sequential entry of a set of 3 different arms. OB^GHSR−/−^ mice showed significantly less spontaneous alternations (Two-tailed unpaired t tests, P=0.0034). **G)** Spontaneous alternation behaviour score, SAB, is the number of alternations divided by alternation opportunities minus 1. The SAB score was not significantly different (Two-tailed unpaired t tests, P=0.0773). **H-K)** A schematic of the Novel arm exploration task created with BioRender.com, wherein in the initial trial one arm is blocked, and after a break, the mice are allowed to explore all arms of the maze in the second trial (WT=10, OB^GHSR−/−^=9). **I)** Novel arm entry: number of entries of the previously blocked arm. **J)** Novel arm duration or time spent in novel arm. **K)** Distance moved. Although, OB^GHSR−/−^ mice had fewer novel arm entries and spent less time in the novel arm, they also moved less in the Y maze (Two-tailed unpaired t tests, *Suppl. Table 1*). **L-O)** A schematic of the Baited Y maze test created with BioRender.com, which measures the ability of the mouse to remember the baited arm (peanut butter chip) in 2x 2-minute training sessions. In the test, the exploration of the previously baited arm was measured (WT=10, OB^GHSR−/−^=9). OB^GHSR−/−^ mice entered the previously baited arm with similar latency (M), spent a similar amount of time in that arm (N) (Two-tailed unpaired t tests, *Suppl. Table 1*) and exhibited similar preference scores as controls in fasted conditions (Two-way ANOVA, P_test_=0.9976, *Suppl. Table 1*).

To further assess working memory as well as reference spatial memory, we used a RAM protocol. In this task, OB^GHSR−/−^ mice also explored the baited arms of the RAM less, and had fewer transitions (**Fig. 7A-D**), but showed no difference in working memory error or calculated memory score (**Fig. 7G,H**) based on the number of re-entries into an already visited arm (working memory error). To test reference spatial memory, we used a delayed win shift protocol (**Fig. 7I**), where 4 arms were blocked and only 4 arms accessible and baited during a training period. The reference error is the number of entries into an arm that was baited in a training period but is not baited during the test. Mice conducted a total of 14 trials within 3 weeks, and consecutive trials within one week were summarized (**Fig. 7J**). Compared to WT, OB^GHSR−/−^ mice needed more time to enter all correct arms and made fewer absolute working memory errors or reference errors but had fewer total arm entries to complete the task, (**Fig. 7K-M**) resulting in no difference in memory score (**Fig. 7N**). In summary, these data support previous findings that OB^GHSR^ are important for exploration, as OB^GHSR−/−^ mice explore less in novel arm of the Y maze, but OB^GHSR^ deletion does not impact on spatial working memory or learning.

**Figure 7.**
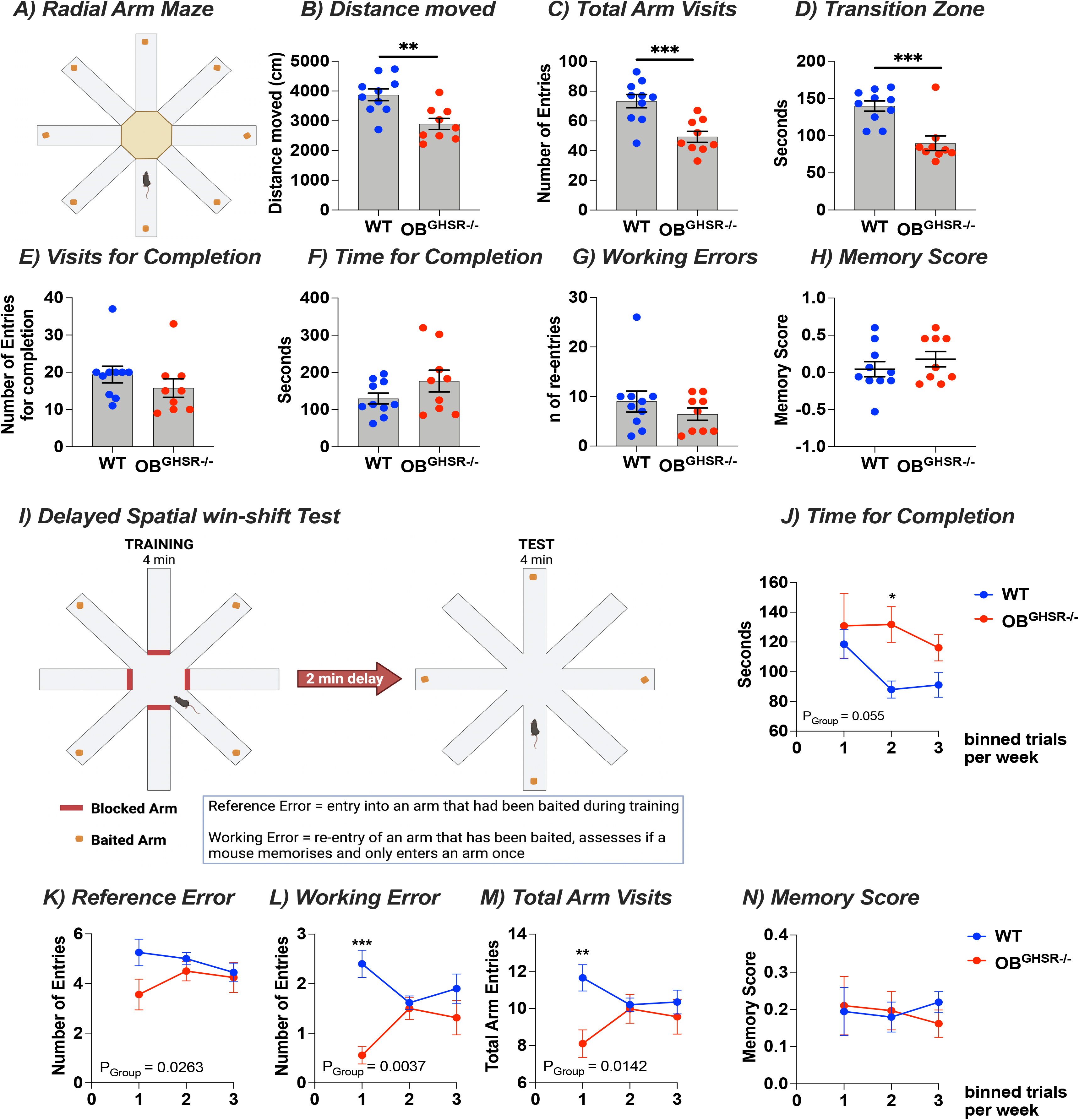
OB GHSR deletion does not affect learning and memory. **A-H)** A schematic of Radial arm maze exploration with 8 baited arms created with BioRender.com, which measures spatial learning and memory (WT=10, OB^GHSR−/−^=9). OB^GHSR−/−^ mice explored the baited arms of the radial arm maze less, as reflected by less distance moved (B) (two-tailed unpaired t tests, P=0.0024), fewer arm visits (C) (two-tailed unpaired t tests, P=0.0008), or time spent in the middle transition zone (D) (two-tailed unpaired t tests, P=0.0005). **E)** Visits for completion. **F)** Time for completion. There was no difference in number of arm visits (E) or time (F) needed to complete the task to visit all 8 arms (two-tailed unpaired t tests, *Suppl. Table 1*). **G)** Working errors: number of re-entries into an already visited arm. **H)** Memory score: the difference of correct and incorrect arm entries divided by the total arm entries. There was no difference in working errors (G) or memory score (H) (two-tailed unpaired t tests, *Suppl. Table 1*). **I-N)** A schematic demonstrating the Delayed win shift test in a radial arm maze, created with BioRender.com, to test learning and memory. Mice are trained that baited arms are alternated during the training and the test phase, while 4 arms are rewarded and the other 4 arms are blocked during the training period. During the test period, all arms are accessible, and the reward is shifted to different arms (WT=10, OB^GHSR−/−^=9). **J)** Time for completion with consecutive trials within one week was summarized. OB^GHSR−/−^ mice needed more time to enter all correct arms (Two-way ANOVA, P_genotype_=0.055, *Suppl. Table 1*). **K)** Reference error (Two-way ANOVA, P_genotype_=0.0263, *Suppl. Table 1*). **L)** Working error (Two-way ANOVA, P_genotype_=0.0037, *Suppl. Table 1*). **M)** Total arm visits (Two-way ANOVA, P_genotype_=0.0142, *Suppl. Table 1*). **N)** Memory score: the difference of correct to incorrect arm entries divided by total arm entries. Although OB^GHSR−/−^ mice made fewer reference (K) or working errors (L), but had fewer total arm entries (M), this resulted in no difference in the calculated memory score (Two-way ANOVA, P_genotype_=0.8391, *Suppl. Table 1*).

### OB^GHSR^ deletion affects body weight and substrate utilisation

Unexpectedly, OB^GHSR−/−^ mice were significantly heavier than control mice, with greater fat mass in the ad libitum or fasted state (**Fig. 8A-D**). There was no difference in the fat excretion in the faeces between the groups (**Suppl. Fig. 7D-F**), showing that differences in fat absorption cannot explain body weight gain in OB^GHSR−/−^ mice. Daily energy expenditure or the amount of energy burnt per hour depends on resting metabolic rate, the thermic effect of food intake, and the energy cost of physical activity. OB^GHSR^ deletion did not change energy expenditure in either the dark or light phase (**Fig. 8E,F**) nor locomotor activity (**Fig. 8I,J**). Interestingly, the absence of GHSRs in the OB resulted in an increase in the respiratory exchange ratio (RER) that was more pronounced in the light phase (**Fig. 8G,H**). The RER is an estimate of which macronutrient is being metabolized for energy, while an RER ratio closer to 1 indicates carbohydrates being metabolized and lipids with a ratio closer to 0.7. Comparison of the RER between the two genotypes shows a significant increase in RER during the light phase in OB^GHSR−/−^ mice, suggesting that in the absence of GHSR signalling in the OB shifts fuel utilization towards carbohydrates whilst promoting fat storage.

**Figure 8.**
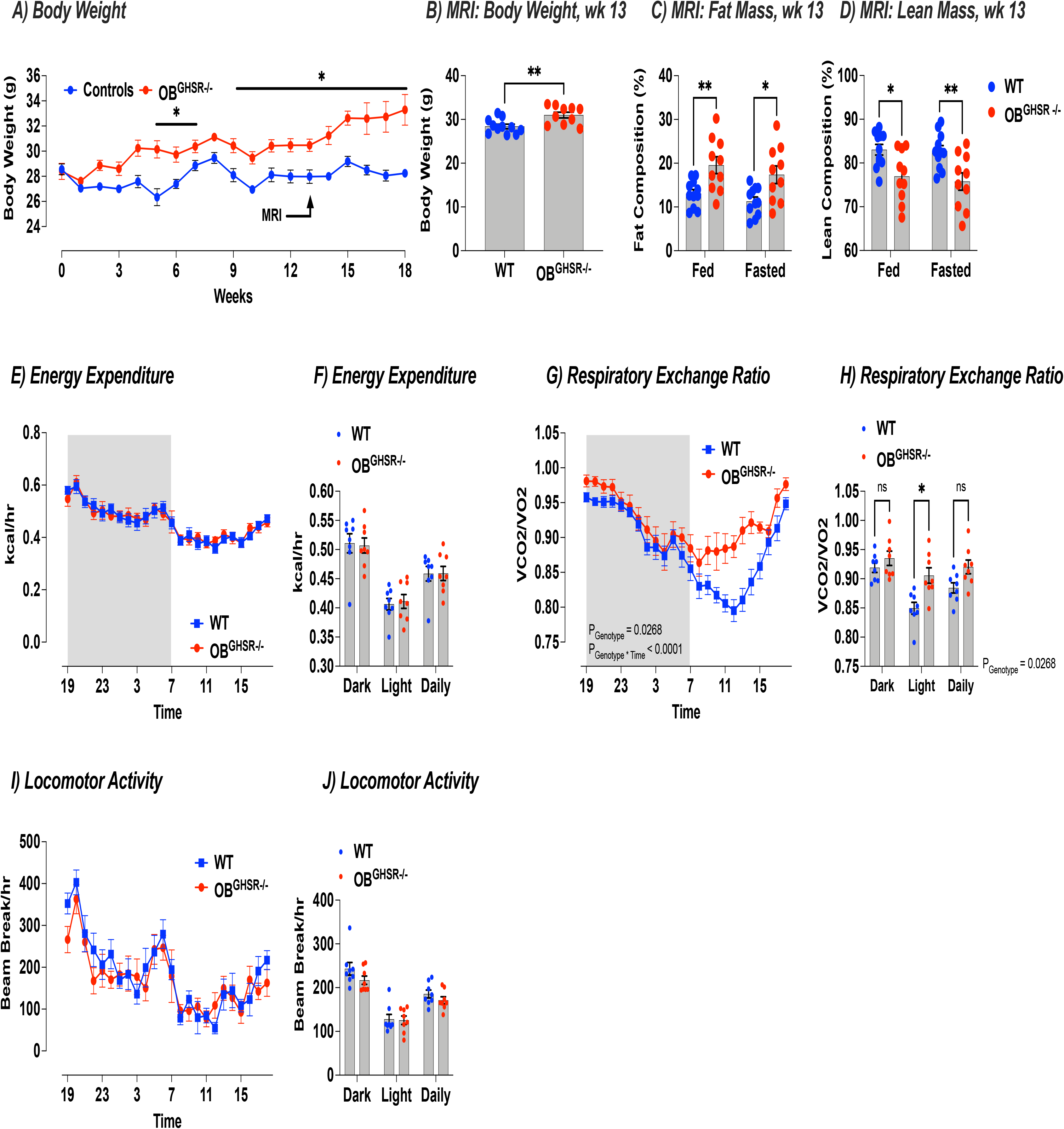
OB GHSR deletion affects energy metabolism. **A)** Body weight (WT= 36, OB^GHSR−/−^=33, 3 cohorts compiled). OB^GHSR−/−^ mice were significantly heavier (Two-way ANOVA, P_genotype_ <0.0001). **B-D)** EchoMRI body composition at 13 weeks post cre injection (WT=11, OB^GHSR−/−^=10). OB^GHSR−/−^ mice exhibited a higher body weight (B) (two-tailed unpaired t test, P=0.0052), with greater fat mass (C) and smaller lean mass (D) in an ad libitum or fasted state (Two-way ANOVA, P_genotype_=0.0002, and P_genotype_=0.0002 respectively, *Suppl. Table 1*). E,F) Energy expenditure (WT= 8, OB^GHSR−/−^=8). OB GHSR deletion did not change energy expenditure in either the dark or light phase (Two-way ANOVA, P_genotype_ =0.9785). G,H) Respiratory exchange ratio. OB^GHSR−/−^ mice displayed an increase in the respiratory exchange ratio that was more pronounced during the light period (Two-way ANOVA, P_genotype_ =0.0268, P_genotyoe*time_ =0.0002, *Suppl. Table 1*). I,J) Locomotor activity. OB GHSR deletion did not change locomotor activity in either the dark or light phase (Two-way ANOVA, P_genotype_ =0.2616).

### OB^GHSR^ deletion affects glucose metabolism

In addition, OB^GHSR^ deletion resulted in impaired glucose metabolism, as OB^GHSR−/−^ mice had higher fasted glucose and insulin levels after a short fast of 4 hours and during a 24-h fasting time course experiment (**Fig. 9A-C**). Fasting corticosterone levels were not increased in OB^GHSR−/−^ mice but tended to be increased after the extended fast of 24h (**Fig. 9E**). Upon refeeding insulin remained elevated with an attenuated effect of refeeding to suppress NEFA (**Fig. 9F-H**). To further evaluate glucose metabolism, we assess glucose tolerance and insulin sensitivity. After an oral administered glucose challenge (2g/kg), glucose clearance was impaired in OB^GHSR^ deleted mice (**Fig. 9I,J**) due to significantly reduced plasma insulin at 15 min during the GTT (**Fig. 9K-M**). During an ITT, insulin administration was less effective to lower blood glucose or NEFA in OB^GHSR−/−^ mice compared to WT controls (**Fig. 9N-O**), suggesting impaired insulin sensitivity in OB^GHSR^ deleted mice. When challenged with 2DG to assess a counterregulatory response to glucopenia, blood glucose was significantly higher in OB^GHSR−/−^ mice (**Fig. 9P,Q**). Collectively, these studies suggest OB^GHSR^ regulate blood glucose independent from changes in body weight since GTT AUC was not correlated with body weight in either WT or KO mice (**Suppl. Fig. 7D**). Since olfactory detection of food can impact on gastric emptying as well as digestion ^15, 42, 43^, we examined gastric emptying during an oral GTT and observed OB^GHSR−/−^ mice had a delayed gastric emptying (**Suppl. Fig. 7E,F**). All this suggests that OB^GHSR^ deletion impaired glucose tolerance, indicated by increased fasted blood glucose and insulin and tested by oral glucose tolerance test (GTT) and insulin tolerance test (ITT). This was independent to differences in body weight, gastric emptying during the GTT or fat absorption (**Suppl. Fig. 7**).

**Figure 9.**
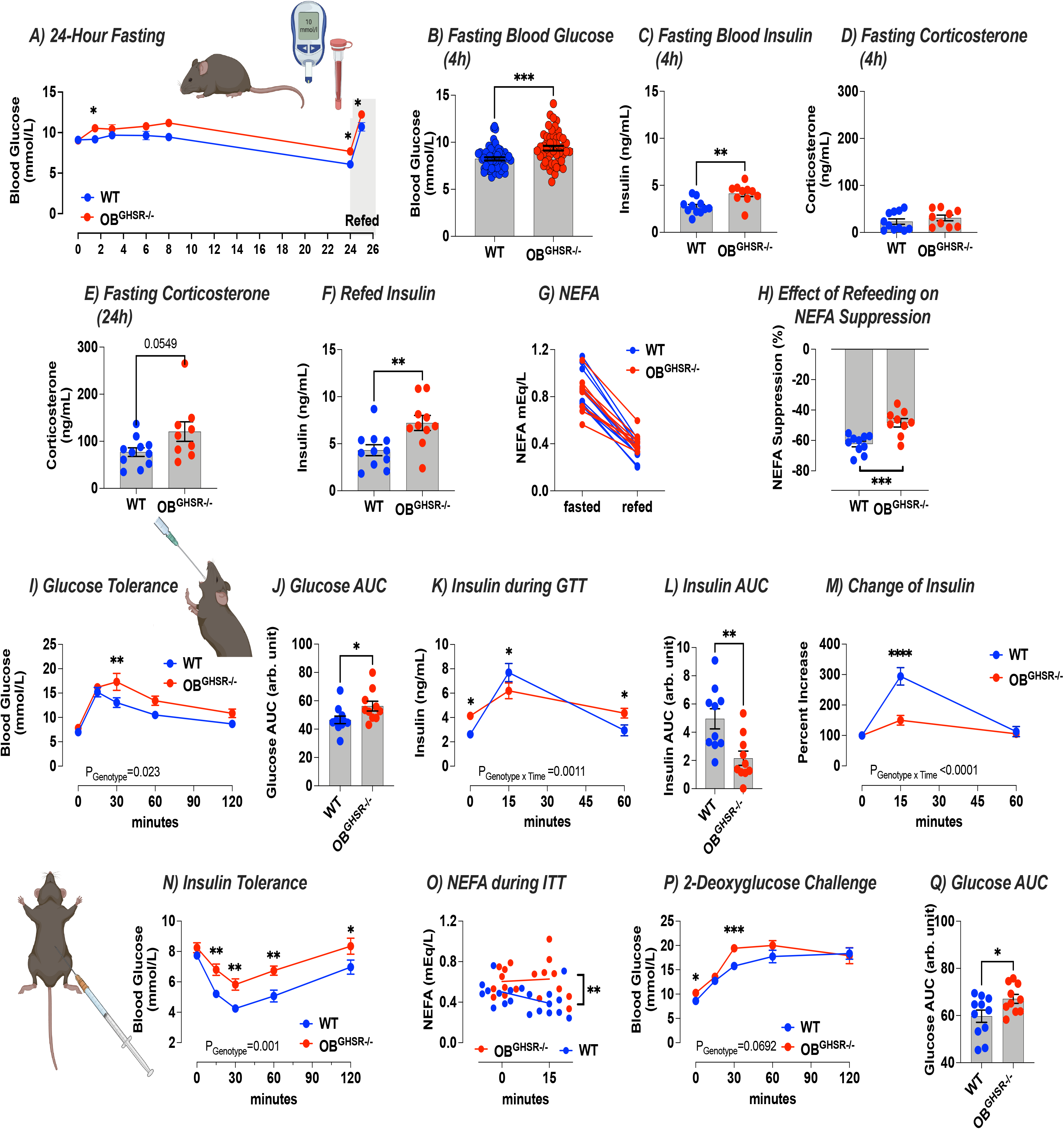
OB GHSR deletion affects glucose metabolism. **A-H)** 24-hour fasting time course experiment (WT= 11, OB^GHSR−/−^=10). Fasting resulted in higher blood glucose in OB^GHSR−/−^ mice during the time course experiment (Two-way ANOVA, P_genotype_ = 0.0025, *Suppl. Table 1*). OB^GHSR−/−^ mice exhibited higher blood glucose (B) and blood insulin levels (C) after 4 hour fasting (Two-tailed unpaired t tests, P=0.0002 and P=0.0023, *Suppl. Table 1*). **D,E)** Blood corticosterone after 4 hour (D), and 24 fasting (E) (Two-tailed unpaired t tests, P=0.369 and P=0.0549). **F-H)** Refeeding with blood insulin (F) and non-esterified free fatty acids (G) after 1 hour of reintroducing food. OB^GHSR−/−^ mice displayed higher blood insulin levels after refeeding (Two-tailed unpaired t tests, P=0.0077). There was no difference between the genotypes for non-esterified free fatty acids (NEFA) levels (Two-way ANOVA, P_metabolic state_ <0.0001, *Suppl. Table 1*). **H)** Effect of refeeding on NEFA suppression (Two-tailed unpaired t tests, P=0.0006). **I-M)** Glucose tolerance test (WT= 11, OB^GHSR−/−^=10). **I)** Blood glucose levels after an oral administration of glucose (2g/kg). OB GHSR deletion impaired glucose clearance (Two-way ANOVA, P_genotype_ =0.0226, *Suppl. Table 1*). **J)** Area under the curve (AUC) of glucose clearance (Two-tailed unpaired t test, P=0.0361). **K)** Blood insulin levels during the glucose tolerance test (Two-way ANOVA, P_genotype*time_ =0.011, *Suppl. Table 1*). **L)** Insulin area under the curve (Two-tailed unpaired t tests, P=0.0049). **M)** Change of insulin during the glucose tolerance test (Two-way ANOVA, P_genotype*time_ <0.0001, *Suppl. Table 1*). **N,O)** Insulin tolerance test (WT= 10, OB^GHSR−/−^=8). **N)** Blood glucose levels after an intraperitoneal injection of insulin (0.75 mU/g). **O)** NEFA. Compared to WT animals, insulin was less effective to lower blood glucose levels, or NEFA (Two-way ANOVA, P_genotype_ =0.001 and P_genotype_ =0.0067, respectively, *Suppl. Table 1*). **P,Q)** 2-Deoxyglucose challenge (WT= 11, OB^GHSR−/−^=10). **P)** Blood glucose levels after an intraperitoneal injection of 2-deoxyglucose (0.5μg/g) (Two-way ANOVA, P_genotype_ =0.0692, *Suppl. Table 1*). **Q)** AUC of glucose during 2-deoxyglucose challenge (Two-tailed unpaired t test, P=0.0369).

## Discussion

Food odours are important sensory cues that convey environmental information about food availability, palatability and calorie content to an organism. These cues invigorate food-seeking, spatial navigation, taste and reward value processing ^12^, as well provide preparatory signals to prime the appropriate metabolic response to incoming nutrients ^1, 9^. With this in mind, it is not surprising that hunger increases olfactory sensitivity ^20, 21, 22, 23^. Here we examined the hypothesis that OB^GHSR^ expression is a novel hunger-signalling mechanism that links metabolic state to olfaction and regulates the behavioural and metabolic consequences of heightened olfactory function. To do this, we deleted OB^GHSRs^ in adult mice with stereotaxic delivery of AAV-cre recombinase using a previously validated *Ghsr* floxed mouse line ^47^ and discovered OB^GHSR^ deletion decreased olfactory sensitivity to food and non-food odours in various olfactory performance tasks in both fed and fasted conditions. Although daily total food intake or ghrelin-induced food intake were not different, OB^GHSR^ deletion decreased the number of feeding bouts in the dark phase and impaired food finding in a buried food seeking test. OB^GHSR^ deletion surprisingly increased body weight, fat mass accumulation by reducing lipid utilisation and impaired glucose tolerance. Moreover, whilst OB^GHSR^ deletion did not affect spatial navigation, it increased anxiety-like behaviour, reduced exploratory behaviour in novel environments, particularly in the fasted state, and increased anhedonia, as highlighted by a reduced preference for 0.1% saccharin. Our results demonstrate a novel role for OB^GHSRs^ in the control of olfactory sensitivity, anxiety-like, exploratory, feeding behaviour (not food consumption) and peripheral energy and glucose metabolism.

Even though the OB has significant GHSR expression ^25, 36, 37^, and the highest uptake of ghrelin in the entire brain ^39^, this is the first study to examine the functional role of GHSRs in the OB. Of note, OB^GHSR^ deletion significantly impaired olfactory sensitivity to palatable food (peanut butter, froot loop), non-food odours (rosewater) and pheromone (urine) odours, as judged by olfactory habituation and olfactory preference tests. Food/non-food and pheromone odours are processed through distinct regions of the OB, with food/non-food odours processed through main olfactory bulb (MOB) and pheromones processed through the accessory olfactory bulb (AOB) ^48, 49^. A deficit in both food/non-food and pheromone detection after OB^GHSR^ deletion is supported by post-mortem expression of cre expression and cre-dependent RFP expression in both the MOB and AOB. This highlights the broad impact and necessity for OB^GHSR^ in normal olfactory function through both the MOB and AOB.

Although olfactory function was impaired in both fed and fasted mice after OB^GHSR^ deletion, fasting was more often associated with greater interest in food odours, such as time spent sniffing froot loops vs rose water and time spent sniffing PB-scented open field. These results are consistent with the idea that fasting increases sensitivity to food odours ^22, 50, 51, 52^, an effect consistently seen across species including humans, mice and flies, and that hunger dampens rival motivations to prioritise food seeking ^53, 54^. Moreover, many of the strongest behavioural effects observed after OB^GHSR^ deletion occurred in fasted mice, including olfactory habituation and preference tests but also most notably in anxiety-like behavioural and buried food finding tests. Indeed, the metabolic functions of the ghrelin system are potentiated in states of metabolic need, such as fasting, and attenuated in states of metabolic excess, culminating in ghrelin resistance in diet-induced obese mice ^28, 29, 30, 32^. These observations have led to the classification of ghrelin as a survival hormone ^29^. The impact of OB^GHSR^ deletion on anxiety-like and anhedonia mirrors the effects of impaired olfactory function on depression and anxiety in humans ^55^ and mice ^56^, as well as the loss of pleasure and enjoyment of food in anosmic humans ^12, 57^. These findings all underscore the use of olfactory bulbectomy as a rodent model of depression-like behaviour ^13^. Interestingly, ghrelin and GHSR function are also linked with reduced anxiety-like and depressive-like behaviour ^58, 59, 60, 61^, including following olfactory bulbectomy in mice ^62^, although the main sites of action have not been fully elucidated. Previous studies suggested the exaggerated depressive-like symptoms associated with *Ghsr* deletion in mice could be overcome by re-expressing GHSRs selectively in catecholaminergic neurons ^63^ or by augmenting hippocampal neurogenesis ^64^, implicating the ventral tegmental area and the hippocampus. Our studies, however, highlight an important and unappreciated role for OB^GHSR^ signaling to alleviate anxiety- depressive-like behaviour, as well exploratory behaviour, particularly in the fasted state. Therefore, our results are consistent with OB^GHSR^ signaling acting to link hunger with olfactory sensitivity to facilitate behavioural adaptation and optimise food seeking and foraging behaviour. Notably, the impairments in anxiety-like behaviour and exploration after OB^GHSR^ deletion were repeated with Y maze and RAM performance, but no deficits in general locomotor activity in metabolic cages or with daily home cage running wheels were observed. These data reinforce the idea the OB^GHSRs^ play a specific anxiolytic role and encourage locomotor exploration to optimise food seeking and foraging.

GHSR deletion in the OB did not affect daily cumulative or daily average food consumption, nor did it affect refeeding after fasting or in response to IP ghrelin. These results are not surprising considering GHSR expression remained intact elsewhere in the brain and body. For example, a number of studies have shown crucial roles for AgRP neurons or TH neurons in the neural control of food intake ^63, 65, 66, 67^. Nevertheless, GHSRs in the OB were crucial for food seeking in a buried-food finding test in short-term fasting, but not fed mice, and in response to IP ghrelin. In addition, feeding behaviour as assessed by bout number and duration, was significantly impaired after OB^GHSR^ deletion. Given the impact of OB^GHSR^ deletion on anxiety-like and exploratory behaviour, we argue that this behaviour phenotype is likely a contributing factor to the impaired food-seeking and feeding behaviour observed. Moreover, this supports the hypothesis that OB^GHSRs^ are required to optimise foraging by facilitating the appropriate behavioural adaptations to low food availability. Hunger-sensing AgRP neurons enhance food odour attraction over pheromone odours, as well as promote food intake ^51^. Although GHSRs are co-expressed on >90% of NPY/AgRP neurons in the ARC ^68, 69^, our discovery demonstrates a specific role for GHSRs in the OB, independent from AgRP neurons in foraging and anxiety-like behaviour but not food consumption. Future studies should examine the impact of GHSR deletion from AgRP neurons on olfactory performance.

Hunger-sensing AgRP neurons are also thought to optimise foraging and recent studies highlight the importance of sensory detection of calorie availability to AgRP neurons ^70^. For example, the sensory detection of food resets activity of hypothalamic AgRP and POMC neuronal activity within seconds, with a response magnitude that predicts the calorie content of the food to be consumed and the energy need of the animal ^2^. Intriguingly, these learned sensory-driven activity changes occur before food consumption but require both metabolic sensing in AgRP neurons ^6^ and post-ingestive calorie feedback from the gut to sustain sensory-driven changes in activity ^71, 72, 73^. These studies suggest that the neural circuits controlling energy balance coordinate sensory input and metabolic feedback over multiple scales ^74^. Moreover, hunger sensing AgRP neurons integrate sensory information related to external food availability and calorie content with the interoceptive energy need of the organism. Olfaction is likely to be a critical system relaying external sensory information of energy availability to AgRP neurons to facilitate this integration. Olfaction is usually the first sensory modality predicting food characteristics and plays an important role in food-seeking for many animals ^7^, and olfactory detection of food odours can acutely, but transiently, suppress AgRP neuronal activity ^51^. Thus, our data predicts the coordinated action of GHSR-dependent hunger-signalling in the OB, together with hunger sensing in hypothalamic AgRP neurons, is required for appropriate behavioural and metabolic response to fasting and low energy availability, although further studies are required to explore this interaction.

While total caloric consumption was not affected by OB^GHSR^ deletion, we observed an increase in body mass and fat mass. This is somewhat surprising given that whole body deletion of the GHSR results in reduced body weight after HFD feeding ^75^, ghrelin deletion protects against early onset of obesity ^76^ and rebound weight gain after diet-induced weight loss ^33^, although not all studies have reported similar findings ^77, 78^. However, olfactory dysfunction is associated with weight gain and obesity ^19, 79, 80, 81^ and increased olfactory sensitivity prevented diet-induced obesity in both genetic pharmacological models ^17, 18, 19^. These studies suggest that weight gain after OB^GHSR^ deletion may be a secondary consequence caused by impaired olfactory sensitivity. The regulation of peripheral lipid utilisation is a likely link since exposure to a food odour triggers fat mobilisation and utilisation in mice ^9^ and worms ^3^. Consistent with this, we observed less fat utilisation and greater carbohydrate utilisation after OB^GHSR^ deletion. Notably, differences in nutrient partitioning are associated with perturbations in body weight gain ^82^.

Deletion of GHSR signalling in the OB also impacted glucose metabolism. OB^GHSR−/−^ mice had higher fasted blood glucose and a limited ability to clear glucose due to impaired insulin secretion, reduced insulin sensitivity and increased glucose production when challenged with 2-deoxyglucose. Our results are in contrast with the observation that conditional ablation of olfactory sensory neurons in the olfactory epithelium prevents insulin resistance and improved glucose clearance ^16^. Nevertheless, olfactory dysfunction is linked to impaired glucose regulation and type 2 diabetes ^83^, in line with our findings.

Our results shed new light on the role of the OB as an integrator of metabolic state and suggest a novel and important neuroendocrine role for the OB. This idea is supported by the expression of various metabolic hormone receptors, including insulin, leptin, cholecystokinin, orexins, and ghrelin ^15, 25, 27^. We hypothesize that olfactory sensitivity is an important sensory integrator of relevant olfactory cues predicting food availability and calorie content. In response to the olfactory detection of known foods or food cues, pre-emptive metabolic and behavioural responses are engaged to facilitate foraging as well as preparation for an incoming meal. This view is informed by the observation that the sensory detection of food or food odours primes hepatic metabolic gene expression and lipid metabolism ^1^. This process was originally described by Pavlov as the “psychic phase of digestion”, when sensory cues, such as food odours, stimulate both salivation and cephalic responses, including to prime the gastrointestinal tract for incoming food (salivation, gastric acid secretion) and insulin release ^7^.

In summary, we show OB^GHSR^ maintains olfactory sensitivity, leading to several behavioural and metabolic adaptations to help an organism respond to low energy availability. We conclude OB^GHSR^ function is a key mechanism linking hunger and energy deprivation with increased olfactory sensitivity and represents a key survival signal that increases sensitivity to salient odour cues in a food-scarce environment. Intact OB^GHSR^ signalling confers appropriate behavioural resilience to explore and exploit foraging opportunities in a potentially anxiogenic environment. At the same time, OB^GHSR^ signalling primes olfactory-driven metabolic responses, including glucose and lipid regulation, thus ensuring appropriate physiological responses to stressful stimuli. An understanding of the precise OB neural circuitry mediating GHSR signalling is an important avenue for future research, especially considering the unique therapeutic potential of intranasal delivery to target pharmacological treatments for metabolic disorders.

## Supporting information

Support Fig 1

Support Fig 2

Support Fig 3

Support Fig 4

Support Fig 5

Support Fig 6

Support Fig 7

## Supplemental Figure Legends

**Suppl. Figure 1 Cre expression**. Cre expression after bilateral stereotaxic injections of AAV-Cre into *Ghsr^fl/fl^*∷Ai14 RFP (OB^GHSR−/−^) and *Ghsr^wt/wt^*∷Ai14 RFP mice (WT) mice.

**Suppl. Figure 2 Olfactory habituation test. A)** Number of behavioural changes in ad libitum fed condition, schematic created with BioRender.com. **B)** Number of behavioural changes in fasted condition. **C)** Number of behavioural changes during fed and fasted conditions. OB^GHSR−/−^ mice exhibited fewer behavioural changes during the olfactory habituation test, suggesting less engagement and behavioural flexibility in exploratory activities (Two-way ANOVA, *Suppl. Table 1*).

**Suppl. Figure 3 Baited open field and female social interaction test. A-F)** A schematic of the Baited open field test created with BioRender.com. **A)** Number of behavioural changes in fed condition. **B)** Behavioural bout frequency in fed condition. **C)** Time spent in a behavioural bout during fed condition. **D)** Number of behavioural changes in fasted condition. **E)** Behavioural bout frequency in fasted condition. **F)** Time spent in a behavioural bout during fasted condition. OB^GHSR−/−^ mice show different behavioural patterns with more stationary behaviour and less involvement in sniffing and walking exploration or behavioural transitions (unpaired t tests, *Suppl. Table 1*). **G-M)** A schematic of a female social interaction test using a 3-chamber preference protocol created with BioRender.com, where mice either explore the 3 chambers with 2 empty wire mesh pencil cups, or a social stimulus (female WT mouse) under one or both of these cups. The amount of time spent interacting with either stimulus was recorded. **H,I)** Object-Object Interaction, 2 empty wire cups. **J,K)** Object-Female Interaction. **L,M)** Female-Female Interaction. No differences in the duration spent in the different zones nor frequencies of investigations between the groups were observed (Two-way ANOVA, *Suppl. Table 1*).

**Suppl. Figure 4 Behavioural anxiety tests. A)** Velocity of elevated plus maze. **B)** Velocity of light-dark box. **C)** Velocity of open field (Two-way ANOVA, *Suppl. Table 1*). Schematics were created with BioRender.com.

**Suppl. Figure 5 Feeding behaviour. A-D)** Feeding behaviour in BioDaq feeding cages (schematic created with BioRender.com; WT= 10, OB^GHSR−/−^=9). **A)** Average Food Intake. **B)** Average Bouts. **C)** Food intake after overnight fasting. **D)** Bouts after overnight fasting. **E)** Food seeking in ad libitum fed conditions. **F)** Food seeking after overnight fasting (Two-tailed unpaired t tests and two-way ANOVAs, *Suppl. Table 1*).

**Suppl. Figure 6 Baited Y Maze in ad libitum fed condition. A)** Baited Y maze protocol created with BioRender.com (WT= 10, OB^GHSR−/−^=9). **B)** Latency. **C)** Percent time spent in previously baited arm. **D)** Distance moved. **E)** Preference Score (Two-tailed unpaired t tests and two-way ANOVAs, *Suppl. Table 1*).

**Suppl. Figure 7 GHSR expression in the olfactory bulb affects metabolism. A)** Faecal triglycerides. **B)** Faecal NEFA. **C)** Faecal NEFA/triglyceride ratio. There was no difference in the fat excretion in the faeces between the groups (WT= 25, OB^GHSR−/−^=21, two-tailed unpaired t tests, *Suppl. Table 1*). **D)** Correlation glucose AUC during GTT with body weight. There was no significant effect of body weight on glucose AUC during the GTT (WT= 11, OB^GHSR−/−^=10, Pearson r correlation, *Suppl. Table 1*). **E)** Gastric emptying rate during the GTT using acetaminophen as a marker for gastric emptying. **F)** AUC of gastric emptying rate. OB^GHSR−/−^ mice had a delayed gastric emptying (WT= 10, OB^GHSR−/−^=10, two-way ANOVA P_genotype_ =0.0456, and two-tailed unpaired t test P=0.0556, *Suppl. Table 1*).

## References

1. Brandt C, et al. Food Perception Primes Hepatic ER Homeostasis via Melanocortin-Dependent Control of mTOR Activation. Cell 175, 1321–1335 e1320 (2018).

2. Chen Y, Lin YC, Kuo TW, Knight ZA. Sensory detection of food rapidly modulates arcuate feeding circuits. Cell 160, 829–841 (2015).

3. Mutlu AS, Gao SM, Zhang H, Wang MC. Olfactory specificity regulates lipid metabolism through neuroendocrine signaling in Caenorhabditis elegans. Nat Commun 11, 1450 (2020).

4. Betley JN, et al. Neurons for hunger and thirst transmit a negative-valence teaching signal. Nature 521, 180–185 (2015).

5. Mandelblat-Cerf Y, et al. Arcuate hypothalamic AgRP and putative POMC neurons show opposite changes in spiking across multiple timescales. Elife 4, (2015).

6. Reichenbach A, et al. Metabolic sensing in AgRP neurons integrates homeostatic state with dopamine signalling in the striatum. Elife 11, (2022).

7. Fine LG, Riera CE. Sense of Smell as the Central Driver of Pavlovian Appetite Behavior in Mammals. Front Physiol 10, 1151 (2019).

8. Jovanovic P, Riera CE. Olfactory system and energy metabolism: a two-way street. Trends Endocrinol Metab 33, 281–291 (2022).

9. Tsuneki H, et al. Food odor perception promotes systemic lipid utilization. Nat Metab, (2022).

10. Wright GA, Mustard JA, Kottcamp SM, Smith BH. Olfactory memory formation and the influence of reward pathway during appetitive learning by honey bees. J Exp Biol 210, 4024–4033 (2007).

11. Croy I, Nordin S, Hummel T. Olfactory Disorders and Quality of Life-An Updated Review. Chemical senses 39, 185–194 (2014).

12. Rolls ET. Taste, olfactory, and food reward value processing in the brain. Prog Neurobiol 127-128, 64–90 (2015)

13. Kelly JP, Wrynn AS, Leonard BE. The olfactory bulbectomized rat as a model of depression: an update. Pharmacol Ther 74, 299–316 (1997).

14. Fernandez-Aranda F, et al. Smell-taste dysfunctions in extreme weight/eating conditions: analysis of hormonal and psychological interactions. Endocrine 51, 256–267 (2016).

15. Palouzier-Paulignan B, et al. Olfaction under metabolic influences. Chemical senses 37, 769–797 (2012).

16. Riera CE, et al. The Sense of Smell Impacts Metabolic Health and Obesity. Cell Metab 26, 198–211 e195 (2017).

17. Fadool DA, et al. Kv1.3 channel gene-targeted deletion produces “Super-Smeller Mice” with altered glomeruli, interacting scaffolding proteins, and biophysics. Neuron 41, 389–404 (2004).

18. Schwartz AB, et al. Olfactory bulb-targeted quantum dot (QD) bioconjugate and Kv1.3 blocking peptide improve metabolic health in obese male mice. J Neurochem 157, 1876–1896 (2021).

19. Tucker K, Overton JM, Fadool DA. Diet-Induced Obesity Resistance of Kv1.3−/− Mice is Olfactory Bulb Dependent. J Neuroendocrinol 24, 1087–1095 (2012).

20. Aime P, Duchamp-Viret P, Chaput MA, Savigner A, Mahfouz M, Julliard AK. Fasting increases and satiation decreases olfactory detection for a neutral odor in rats. Behavioural Brain Research 179, 258–264 (2007).

21. Cameron JD, Goldfield GS, Doucet E. Fasting for 24 h improves nasal chemosensory performance and food palatability in a related manner. Appetite 58, 978–981 (2012)

22. Hanci D, Altun H. Hunger state affects both olfactory abilities and gustatory sensitivity. Eur Arch Otorhinolaryngol 273, 1637–1641 (2016).

23. Jacquot C, Baudoin C. Foraging behavioural changes induced by conspecific and heterosubspecific odours in two strains of wild mice. Behavioural Processes 58, 115–123 (2002).

24. Tiret P, Chaigneau E, Lecoq J, Charpak S. Two-photon imaging of capillary blood flow in olfactory bulb glomeruli. Methods Mol Biol 489, 81–91 (2009).

25. Loch D, Breer H, Strotmann J. Endocrine Modulation of Olfactory Responsiveness: Effects of the Orexigenic Hormone Ghrelin. Chemical senses 40, 469–479 (2015).

26. Martin B, Maudsley S, White CM, Egan JM. Hormones in the naso-oropharynx: endocrine modulation of taste and smell. Trends Endocrinol Metab 20, 163–170 (2009).

27. Tong J, et al. Ghrelin enhances olfactory sensitivity and exploratory sniffing in rodents and humans. Journal of Neuroscience 31, 5841–5846 (2011).

28. Briggs DI, Andrews ZB. Metabolic status regulates ghrelin function on energy homeostasis. Neuroendocrinology 93, 48–57 (2011).

29. Mani BK, Zigman JM. Ghrelin as a Survival Hormone. Trends in Endocrinology and Metabolism 28, 843–854 (2017).

30. Zigman JM, Bouret SG, Andrews ZB. Obesity Impairs the Action of the Neuroendocrine Ghrelin System. Trends Endocrinol Metab 27, 54–63 (2016).

31. Beutler LR, et al. Obesity causes selective and long-lasting desensitization of AgRP neurons to dietary fat. Elife 9, (2020).

32. Briggs DI, Enriori PJ, Lemus MB, Cowley MA, Andrews ZB. Diet-induced obesity causes ghrelin resistance in arcuate NPY/AgRP neurons. Endocrinology 151, 4745–4755 (2010).

33. Briggs DI, Lockie SH, Wu Q, Lemus MB, Stark R, Andrews ZB. Calorie-restricted weight loss reverses high-fat diet-induced ghrelin resistance, which contributes to rebound weight gain in a ghrelin-dependent manner. Endocrinology 154, 709–717 (2013)

34. Mazzone CM, et al. High-fat food biases hypothalamic and mesolimbic expression of consummatory drives. Nat Neurosci 23, 1253–1266 (2020).

35. Zigman JM, Jones JE, Lee CE, Saper CB, Elmquist JK. Expression of ghrelin receptor mRNA in the rat and the mouse brain. Journal of Comparative Neurology 494, 528–548 (2006).

36. Ratcliff M, et al. Calorie restriction activates new adult born olfactory-bulb neurones in a ghrelin-dependent manner but acyl-ghrelin does not enhance subventricular zone neurogenesis. J Neuroendocrinol 31, (2019).

37. Mani BK, et al. Neuroanatomical characterization of a growth hormone secretagogue receptor-green fluorescent protein reporter mouse. The Journal of comparative neurology 522, 3644–3666 (2014).

38. Okuhara Y, Kaiya H, Teraoka H, Kitazawa T. Structural determination, distribution, and physiological actions of ghrelin in the guinea pig. Peptides 99, 70–81 (2018).

39. Diano S, et al. Ghrelin controls hippocampal spine synapse density and memory performance. Nat Neurosci 9, 381–388 (2006).

40. Rhea EM, Salameh TS, Gray S, Niu J, Banks WA, Tong J. Ghrelin transport across the blood-brain barrier can occur independently of the growth hormone secretagogue receptor. Mol Metab 18, 88–96 (2018).

41. Banks WA, Burney BO, Robinson SM. Effects of triglycerides, obesity, and starvation on ghrelin transport across the blood-brain barrier. Peptides 29, 2061–2065 (2008).

42. Shankar K, et al. LEAP2 deletion in mice enhances ghrelin’s actions as an orexigen and growth hormone secretagogue. Mol Metab 53, 101327 (2021).

43. Lopatina O, et al. Anxiety- and depression-like behavior in mice lacking the CD157/BST1 gene, a risk factor for Parkinson’s disease. Front Behav Neurosci 8, 133 (2014).

44. Huang G, et al. Depression-/Anxiety-Like Behavior Alterations in Adult Slit2 Transgenic Mice. Front Behav Neurosci 14, 622257 (2020).

45. Decarie-Spain L, et al. Ventral hippocampus-lateral septum circuitry promotes foraging-related memory. Cell Rep 40, 111402 (2022).

46. Kanoski SE, et al. Hippocampal leptin signaling reduces food intake and modulates food-related memory processing. Neuropsychopharmacology 36, 1859–1870 (2011).

47. Gupta D, et al. Disrupting the ghrelin-growth hormone axis limits ghrelin’s orexigenic but not glucoregulatory actions.. Mol Metab 53, 101258 (2021).

48. Mucignat-Caretta C. The rodent accessory olfactory system. J Comp Physiol A Neuroethol Sens Neural Behav Physiol 196, 767–777 (2010).

49. Xu F, et al. Simultaneous activation of mouse main and accessory olfactory bulbs by odors or pheromones. The Journal of comparative neurology 489, 491–500 (2005).

50. Albrecht J, et al. Olfactory detection thresholds and pleasantness of a food-related and a non-food odour in hunger and satiety. Rhinology 47, 160–165 (2009).

51. Horio N, Liberles SD. Hunger enhances food-odour attraction through a neuropeptide Y spotlight. Nature 592, 262–266 (2021).

52. Sayin S, et al. A Neural Circuit Arbitrates between Persistence and Withdrawal in Hungry Drosophila. Neuron 104, 544–558 e546 (2019).

53. Burnett CJ, et al. Need-based prioritization of behavior. Elife 8, (2019).

54. Sutton AK, Krashes MJ. Integrating Hunger with Rival Motivations. Trends Endocrinol Metab 31, 495–507 (2020).

55. Kohli P, Soler ZM, Nguyen SA, Muus JS, Schlosser RJ. The Association Between Olfaction and Depression: A Systematic Review. Chemical senses 41, 479–486 (2016).

56. Glinka ME, et al. Olfactory Deficits Cause Anxiety-Like Behaviors in Mice. Journal of Neuroscience 32, 6718–6725 (2012).

57. Croy I, et al. Olfaction as a marker for depression in humans. Journal of Affective Disorders 160, 80–86 (2014).

58. Asakawa A, et al. A role of ghrelin in neuroendocrine and behavioral responses to stress in mice. Neuroendocrinology 74, 143–147 (2001).

59. Huang HJ, et al. Ghrelin alleviates anxiety- and depression-like behaviors induced by chronic unpredictable mild stress in rodents. Behav Brain Res 326, 33–43 (2017).

60. Lutter M, et al. The orexigenic hormone ghrelin defends against depressive symptoms of chronic stress. Nat Neurosci 11, 752–753 (2008).

61. Spencer SJ, et al. Ghrelin Regulates the Hypothalamic-Pituitary-Adrenal Axis and Restricts Anxiety After Acute Stress. Biological Psychiatry 72, 457–465 (2012).

62. Carlini VP, et al. Acute ghrelin administration reverses depressive-like behavior induced by bilateral olfactory bulbectomy in mice. Peptides 35, 160–165 (2012).

63. Chuang JC, et al. Ghrelin mediates stress-induced food-reward behavior in mice. J Clin Invest 121, 2684–2692 (2011).

64. Walker AK, et al. The P7C3 class of neuroprotective compounds exerts antidepressant efficacy in mice by increasing hippocampal neurogenesis. Mol Psychiatry 20, 500–508 (2015).

65. Chen HY, et al. Orexigenic action of peripheral ghrelin is mediated by neuropeptide Y and agouti-related protein. Endocrinology 145, 2607–2612 (2004).

66. Luquet S, Phillips CT, Palmiter RD. NPY/AgRP neurons are not essential for feeding responses to glucoprivation. Peptides 28, 214–225 (2007).

67. Wang Q, et al. Arcuate AgRP neurons mediate orexigenic and glucoregulatory actions of ghrelin. Mol Metab 3, 64–72 (2014).

68. Campbell JN, et al. A molecular census of arcuate hypothalamus and median eminence cell types. Nat Neurosci 20, 484–496 (2017).

69. Willesen MG, Kristensen P, Romer J. Co-localization of growth hormone secretagogue receptor and NPY mRNA in the arcuate nucleus of the rat. Neuroendocrinology 70, 306–316 (1999).

70. Reed F, Lockie SH, Reichenbach A, Foldi CJ, Andrews ZB. Appetite to learn: An allostatic role for AgRP neurons in the maintenance of energy balance. Current Opinion in Endocrine and Metabolic Research 24, (2022).

71. Beutler LR, Chen Y, Ahn JS, Lin YC, Essner RA, Knight ZA. Dynamics of Gut-Brain Communication Underlying Hunger. Neuron 96, 461–475 e465 (2017).

72. Goldstein N, McKnight AD, Carty JRE, Arnold M, Betley JN, Alhadeff AL. Hypothalamic detection of macronutrients via multiple gut-brain pathways. Cell Metab 33, 676–687 e675 (2021).

73. Su ZW, Alhadeff AL, Betley JN. Nutritive, Post-ingestive Signals Are the Primary Regulators of AgRP Neuron Activity. Cell Rep 21, 2724–2736 (2017).

74. Chen Y, Knight ZA. Making sense of the sensory regulation of hunger neurons. Bioessays 38, 316–324 (2016).

75. Zigman JM, et al. Mice lacking ghrelin receptors resist the development of diet-induced obesity. The Journal of clinical investigation 115, 3564–3572 (2005).

76. Wortley KE, et al. Absence of ghrelin protects against early-onset obesity. J Clin Invest 115, 3573–3578 (2005)

77. McFarlane MR, Brown MS, Goldstein JL, Zhao TJ. Induced ablation of ghrelin cells in adult mice does not decrease food intake, body weight, or response to high-fat diet. Cell Metab 20, 54–60 (2014).

78. Sun Y, Butte NF, Garcia JM, Smith RG. Characterization of adult ghrelin and ghrelin receptor knockout mice under positive and negative energy balance. Endocrinology 149, 843–850 (2008).

79. Morris A. OBESITY Olfactory senses linked to metabolism. Nature Reviews Endocrinology 13, 494–494 (2017).

80. Peng M, Coutts D, Wang T, Cakmak YO. Systematic review of olfactory shifts related to obesity. Obesity Reviews 20, 325–338 (2019).

81. Vega MV, Rivas AMO. Association of Olfactory Sensitivity with Energy Intake: Role in Development of Obesity. Nutricion Hospitalaria 32, 2385–2389 (2015).

82. Denis RG, et al. Central orchestration of peripheral nutrient partitioning and substrate utilization: implications for the metabolic syndrome. Diabetes Metab 40, 191–197 (2014).

83. Zaghloul H, Pallayova M, Al-Nuaimi O, Hovis KR, Taheri S. Association between diabetes mellitus and olfactory dysfunction: current perspectives and future directions. Diabetic Medicine 35, 41–52 (2018).

