## Supplementary figures and images for "Hunger signalling in the olfactory bulb primes exploration, food-seeking and peripheral metabolism"

### Support Fig 1

*SUPPL. FIGURE 1*

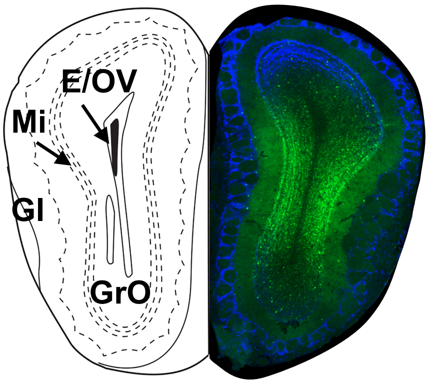


***Bregma 4.28 mm***

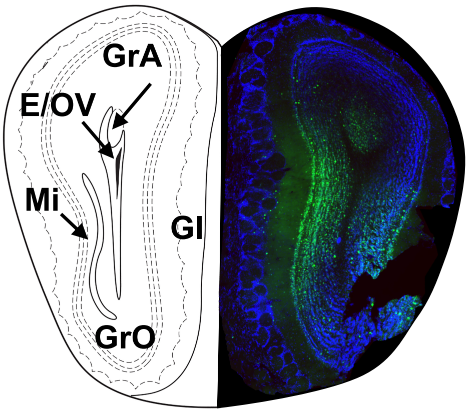


***Bregma 3.92 mm***

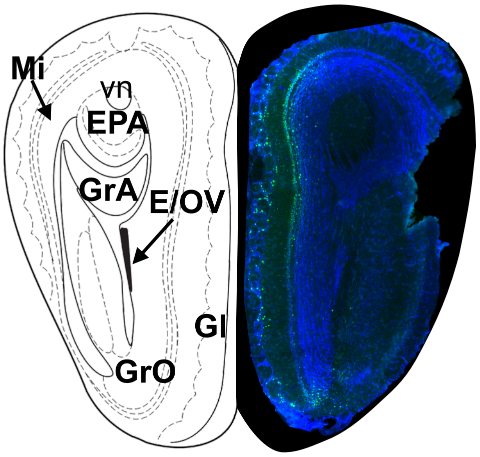


***Bregma 3.56 mm***

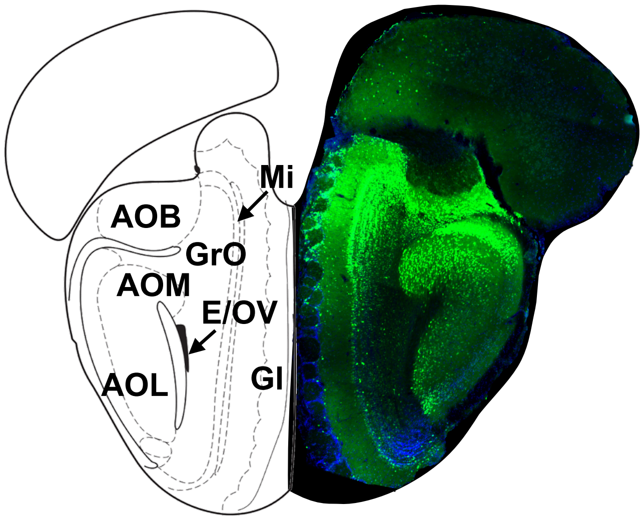


***Bregma 3.2 mm***

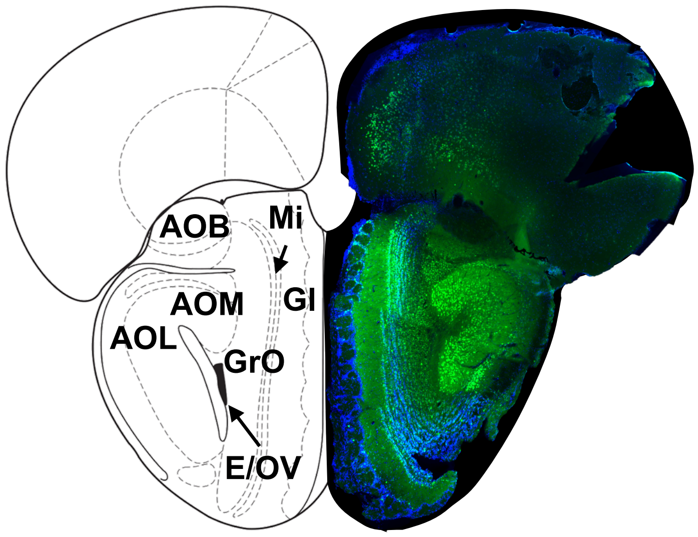


***Bregma 3.08 mm***

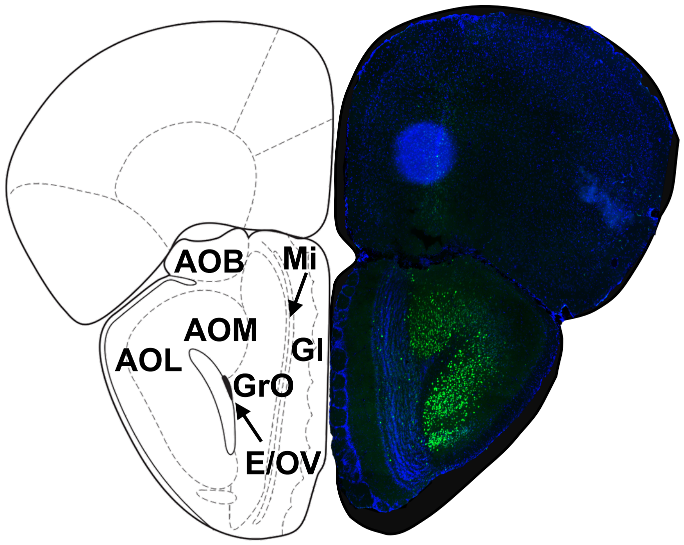


***Bregma 2.96 mm***

### Support Fig 3

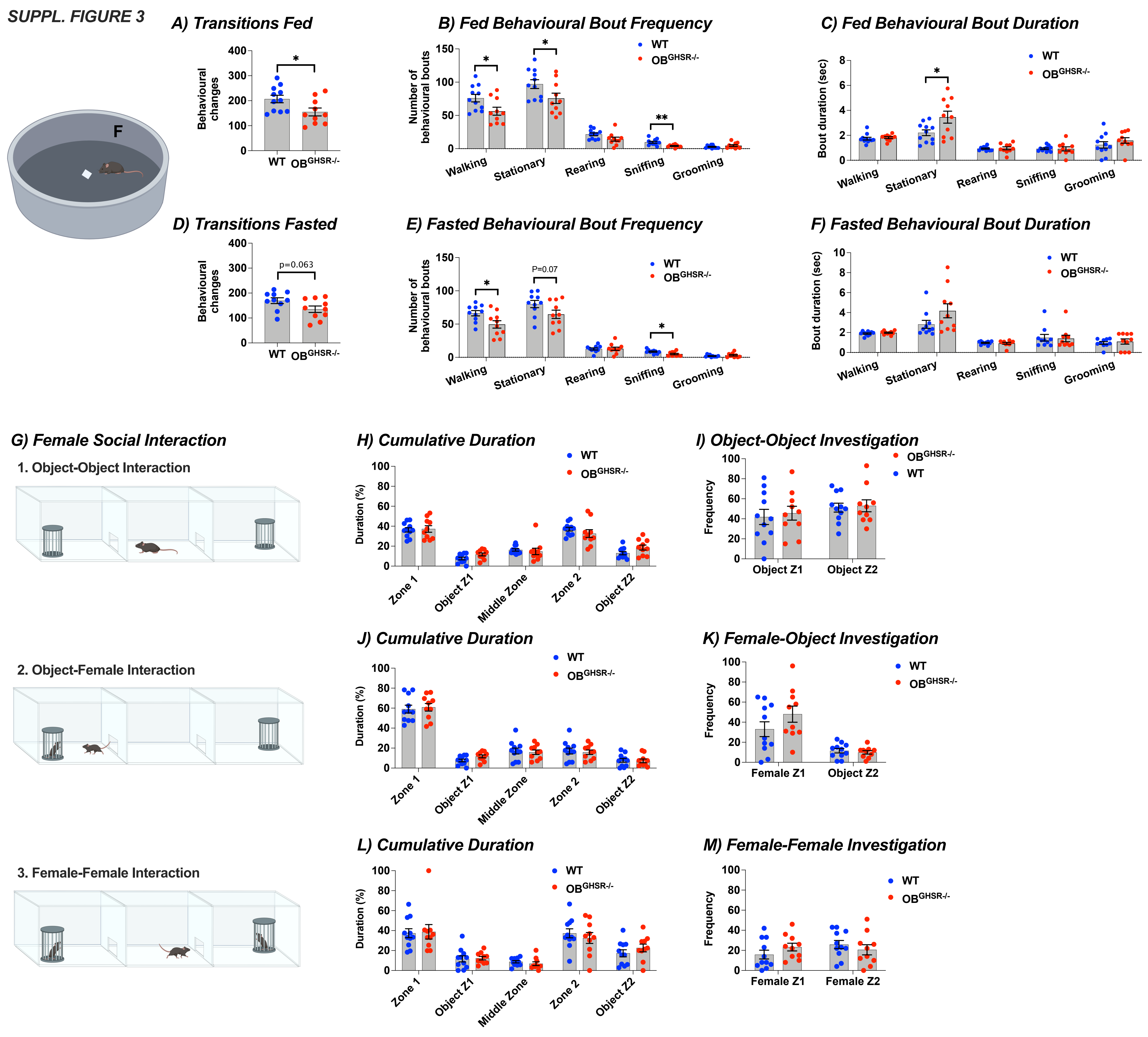

### Support Fig 4

SUPPL. FIGURE 4

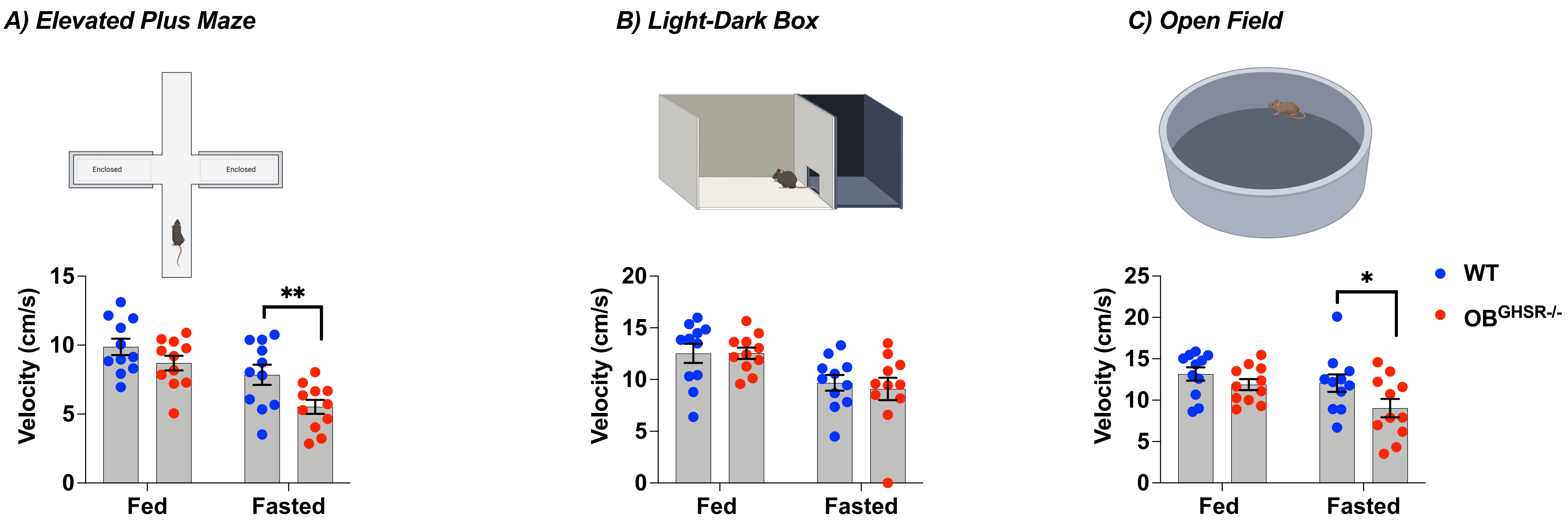

### Support Fig 5

SUPPL. FIGURE 5

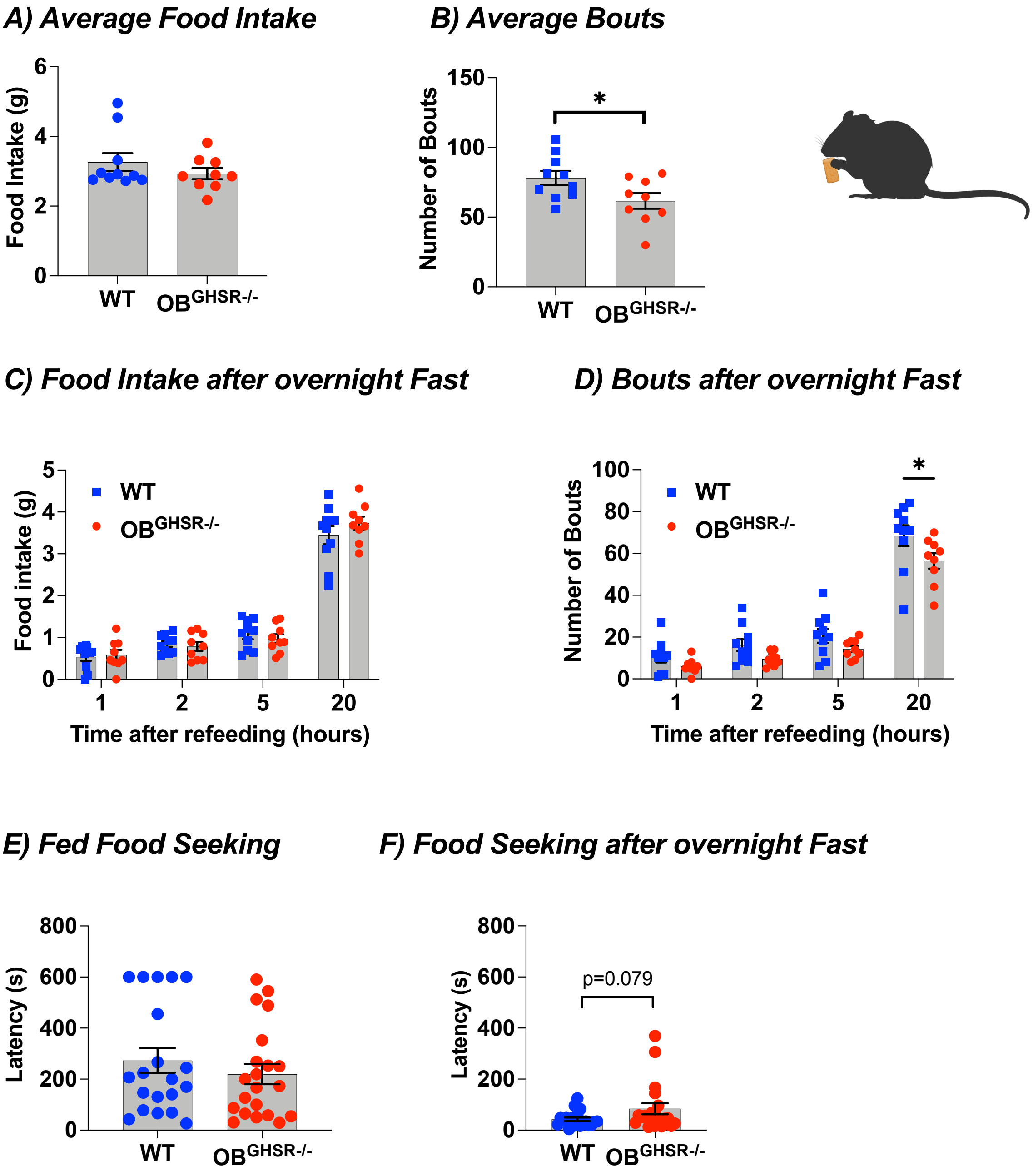

### Support Fig 7

*SUPPL. FIGURE 7*

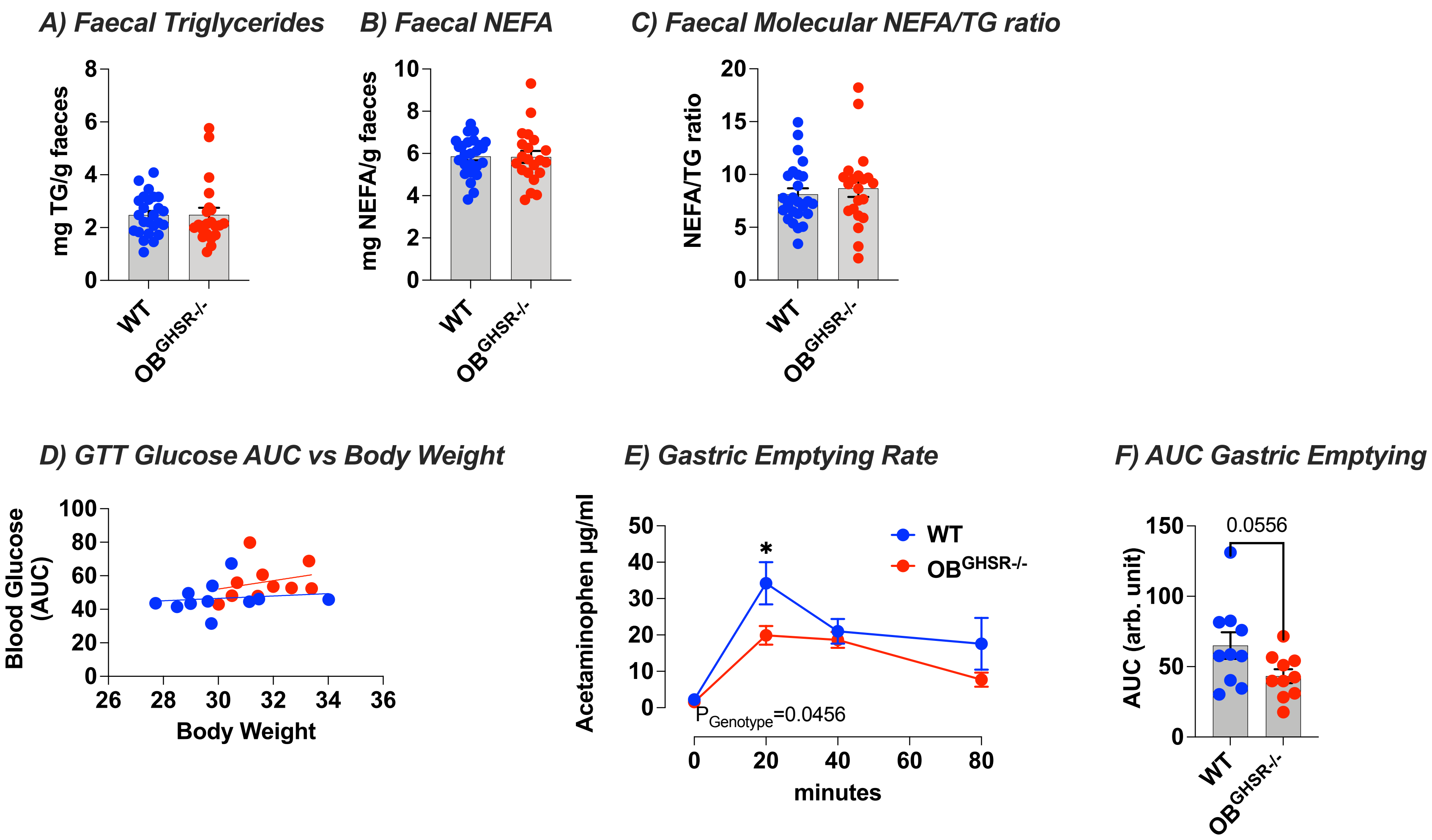
