## Supplementary material for "Hunger signalling in the olfactory bulb primes exploration, food-seeking and peripheral metabolism": Support Fig 2

SUPPL. FIGURE 2

A) Number of Behavioural Changes in the ad libitum Fed state

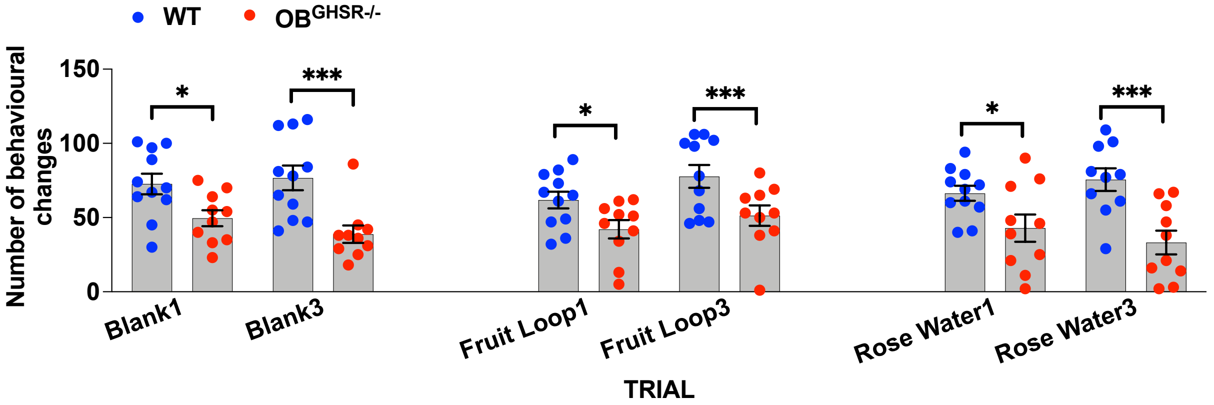


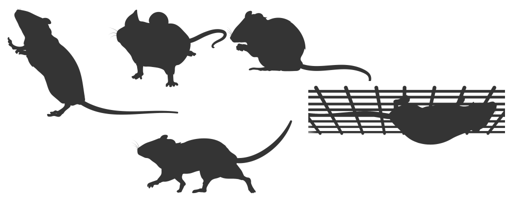

B) Number of Behavioural Changes in the Fasted state

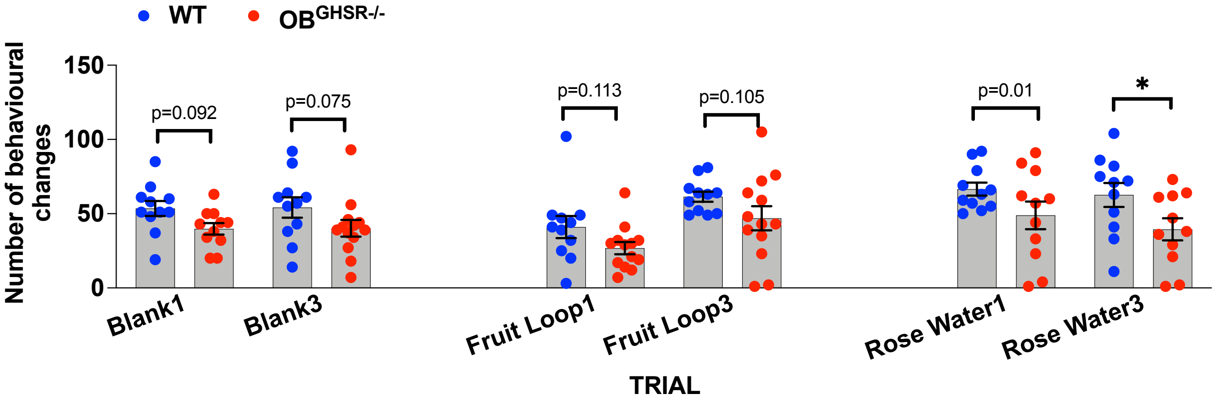


C) All transitions Fed vs Fasted

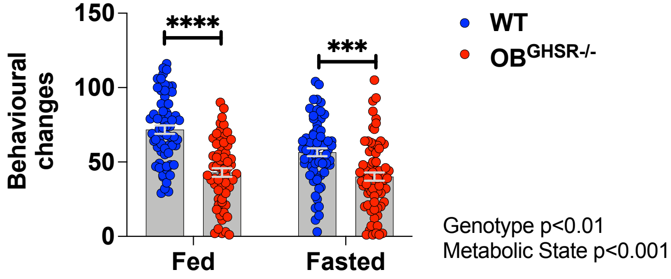
