## Supplementary material for "Hunger signalling in the olfactory bulb primes exploration, food-seeking and peripheral metabolism": Support Fig 6

### *SUPPL. FIGURE 6*

#### *A) Baited Y Maze* *B) Latency* *C) Duration* *D) Distance moved*

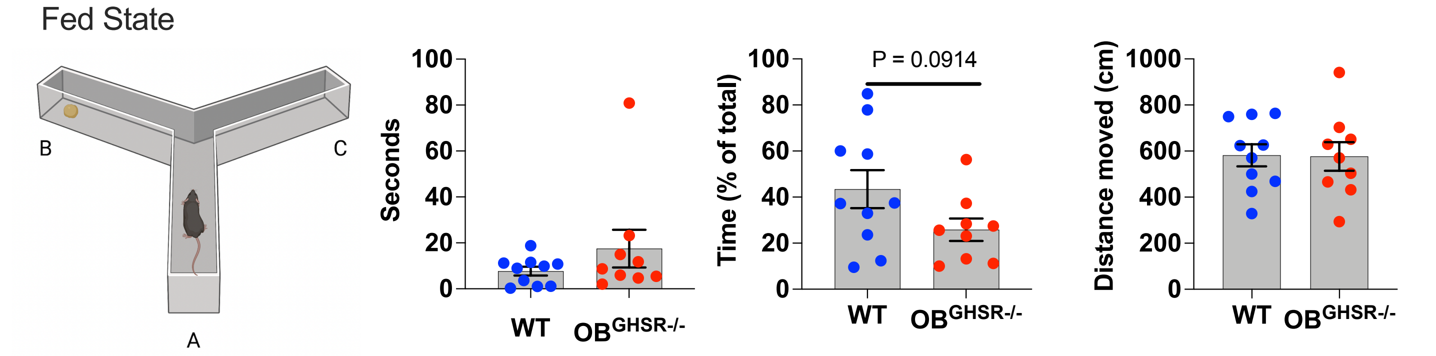


Trial 1 + 2: two training sessions with one baited arm

Trial 3 = TEST: bait removed - previously baited arm visits?

#### *E) Preference Score*

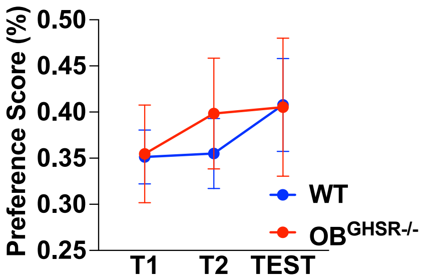
